# GH1-HMGA proteins restrict RNA-directed DNA methylation and antagonize linker histone H1 at euchromatic transposable elements

**DOI:** 10.64898/2026.09.08.749845

**Authors:** Léa Feit, Alejandro Edera, Emmanuel Vanrobays, Lauriane Simon, Delphine Dardalhon-Cuménal, Saad En Naimani, Zdravko J. Lorković, Frederic Berger, Fredy Barneche, Christophe Tatout, Leandro Quadrana, Aline V. Probst, Simon Amiard

## Abstract

In plant genomes, transposable elements (TEs) are silenced by CG and non-CG DNA methylation, with non-CG methylation maintained by CMT chromomethylases and the RNA-directed DNA methylation (RdDM) pathway. In Arabidopsis, linker histone H1 differentially influences these pathways, as loss of H1 causes non-CG hypermethylation at pericentromeric TEs but hypomethylation at euchromatic TEs in gene-rich regions. The molecular basis of this antagonism has remained unclear. Here, we show that GH1-HMGAs, a family of H1-related High Mobility Group proteins, bind AT-rich sequences at euchromatic TEs and prevent RdDM-dependent non-CG methylation by limiting siRNA accumulation. Furthermore, GH1-HMGA and H1 compete for chromatin association at these TEs, as loss of GH1-HMGA triggers ectopic H1 redistribution, while loss of H1 increases GH1-HMGA enrichment, providing a mechanistic explanation for the non-CG hypomethylation observed in *h1* mutants. Collectively, these findings reveal how GH1-HMGA proteins specifically restrain RdDM activity at euchromatic TEs and elucidate how the H1/GH1-HMGA balance spatially partitions distinct DNA methylation pathways across the genome.

## Introduction

Chromatin-based mechanisms are central to the transcriptional repression of transposable elements (TEs), which constitute a major fraction of plant genomes and represent a potent source of genomic instability. In *Arabidopsis thaliana*, TE silencing relies on a self-reinforcing interplay between DNA methylation and histone variants and modifications. DNA methylation occurs in three cytosine sequence contexts, CG, CHG and CHH, maintained by distinct yet interconnected pathways. While MET1 maintains CG methylation, CHROMOMETHYLASE3 (CMT3) and CHROMOMETHYLASE2 (CMT2) are responsible for CHG and CHH methylation, respectively ^1^. Heterochromatic TEs are primarily defined by their pericentromeric localization and high levels of dimethylation of histone H3 at lysine 9 (H3K9me2), which is catalyzed by SUVH-family methyltransferases and recruits CMT2 and CMT3, thereby coupling histone and DNA methylation ^2^. Complementarily, the RNA-directed DNA methylation (RdDM) pathway involves the DRM2 methyltransferase, which is targeted by short interfering RNAs produced by RNA polymerase IV and preferentially acts on smaller, euchromatic TEs interspersed within gene-rich regions ^3^. This spatial partitioning is thought to contribute to the robust control of TE activity in distinct chromatin environments.

Beyond DNA methylation and histone modifications, chromatin structural organization plays a critical role in shaping DNA methylation. One such key determinant is the linker histone H1, which binds the nucleosome dyad and linker DNA *via* its globular H1 (GH1) domain to promote chromatin compaction and constrain TE-associated methylation pathways. In Arabidopsis, loss of H1 results in increased CHH methylation at heterochromatic TEs, largely mediated by CMT2, and concomitantly a reduction in RdDM-dependent methylation at euchromatic TEs ^4,5^. These contrasting effects suggest that H1 acts as a gatekeeper that differentially restricts or permits access of DNA methyltransferases depending on chromatin context, thereby reinforcing the functional distinction between heterochromatic and euchromatic TE silencing pathways. However, the molecular basis underlying H1’s opposing influences on CMT- and RdDM-dependent methylation remains unclear.

In *Arabidopsis thaliana*, the GH1 domain of H1 is also present in other chromatin-associated proteins ^6^, including the GH1-HMGA family. GH1-HMGA1, GH1-HMGA2 and GH1-HMGA3 are partially redundant chromatin proteins that combine a GH1 domain with AT-hook DNA binding motifs and have been implicated in telomere maintenance ^7^ and chromatin looping ^8^. These findings position H1 and GH1-HMGA proteins as chromatin regulators with shared structural features, raising the possibility that they may cooperate or compete at TE chromatin. Yet, whether H1 and GH1-HMGA proteins jointly control TE targeting and DNA methylation remains unknown.

Here, we address this question by profiling the subnuclear and epigenomic distribution of GH1-HMGA and H1 proteins in *Arabidopsis thaliana* wild-type plants and respective mutant lines. Combined with small RNA-seq and genetic depletion of specific DNA methylation machinery components, we further decipher their respective influences on heterochromatic and euchromatic TE methylation. This provided evidence for a previously unrecognized H1/GH1-HMGA homeostatic regulatory layer that differentially modulates TE silencing across chromatin contexts.

## Results

### Evolutionary emergence and diversification of GH1-HMGA proteins with high phase separation potential in dicotyledons

To trace the evolutionary history of the GH1-HMGA protein family, we conducted a phylogenetic analysis of protein sequences with the typical combination of a GH1 domain and AT-hook motifs across representative species of the green lineage **(Figure 1A, Supplementary Figure 1A)**. While mosses and ferns encode H1 proteins, they lack GH1-HMGA proteins, which emerged in angiosperms. In basal angiosperms, we identified four groups of distinct GH1-HMGA-related proteins, yet their architectures differ in both length and number of AT-hook motifs. A group of proteins harbours up to twelve AT-hook motifs, a trait not conserved in other angiosperms. They were therefore classified as "other GH1-HMGA" proteins. In monocotyledons, only small GH1-HMGA proteins (about 200 residues) displaying four AT-hook motifs like Arabidopsis GH1-HMGA3 can be detected. By contrast, dicotyledons share an additional GH1-HMGA paralog, GH1-HMGA2, characterized by a longer size (up to 500 residues), six AT-hook motifs, and an additional A domain of unknown function **(Figure 1B)** ^6^. A third paralog, GH1-HMGA1, emerged alongside the diversification of Brassicaceae. Consequently, GH1-HMGA3 appears to be the ancestral plant GH1-HMGA, while GH1-HMGA1 represents the most recently evolved member of this protein family across angiosperms. GH1-HMGA1 and GH1-HMGA2 share a disordered architecture, with only the GH1 and the A domains being structured. Thermodynamic analysis of standard enthalpy change (ΔH°) using ParSe 2.0 ^9^ as well as Plant PS prediction scores ^10^ served to predict the propensity of GH1-HMGA proteins to form liquid-liquid phase separation (LLPS). Given their capacity to promote phase-separation-mediated heterochromatin aggregation, linker histone sequences were used as a positive reference ^11^. H1.1, H1.2 (ΔH° of 3265 and 3420, respectively), GH1-HMGA1 (2372.6), and GH1-HMGA2 (2100.3) display a high phase separation potential and also fall within the high-confidence Plant PS category (**Figure 1C, Supplementary Figure 1B**). In contrast, GH1-HMGA3 showed only modest autonomous potential (ΔH° = 139.6) and did not meet the Plant PS high-confidence cutoff, consistent with its previously reported ADCP1-dependent function in chromocenter condensation ^12^. Together, these data suggest that GH1-HMGA1 and GH1-HMGA2 proteins have acquired the ability to promote phase separation, whereas GH1-HMGA3, which appears to be the earliest GH1-HMGA member, is predicted to lack this property.

**Figure 1:**
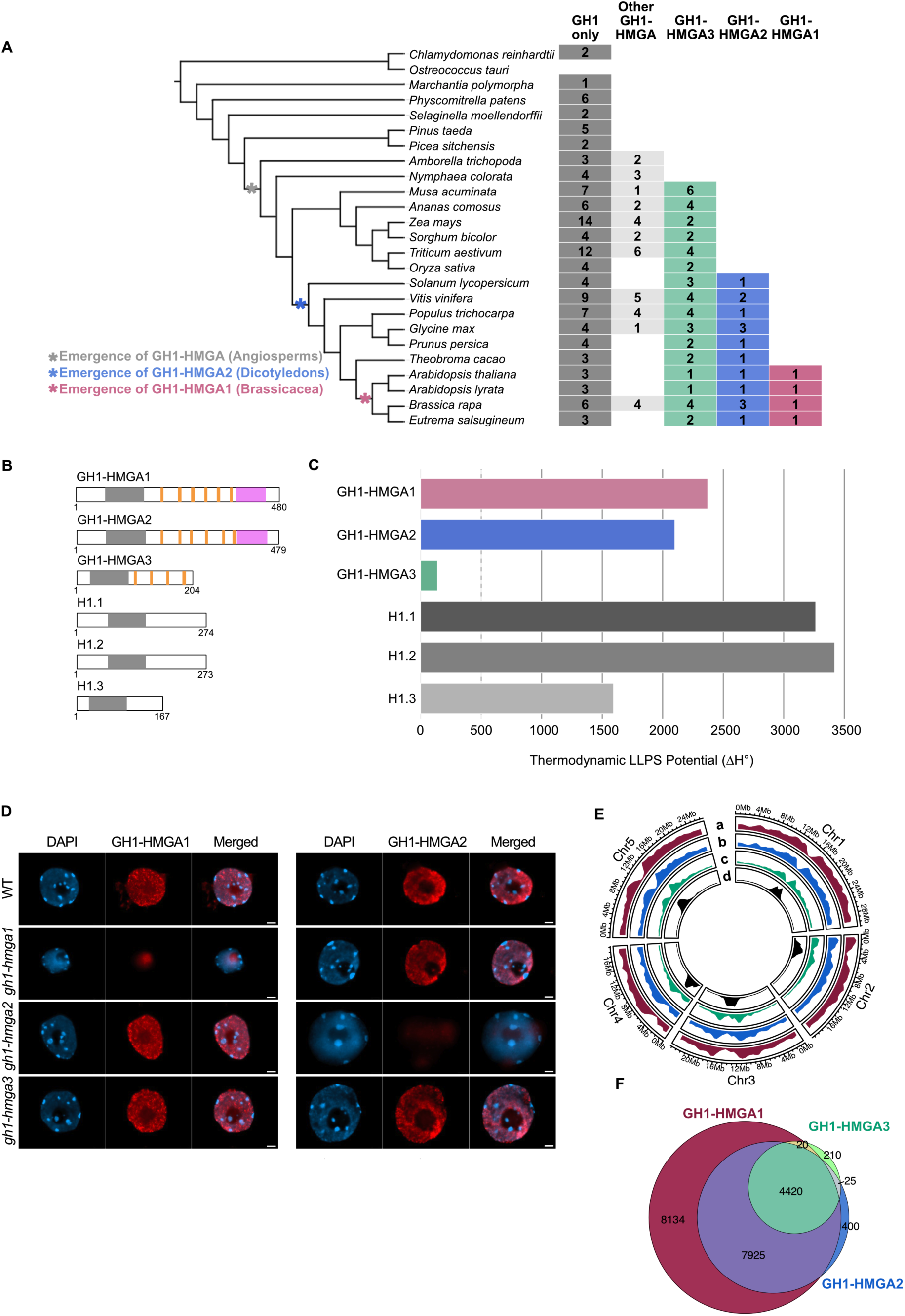
Evolutionary Diversification and Overlapping Target Binding of GH1-HMGA proteins. **(A)** Presence or absence of Arabidopsis H1 protein orthologs (GH1-domain only) and GH1-HMGA proteins (additionally displaying A/T-hook motifs) across plant species spanning the evolutionary history of land plants. GH1-HMGA orthologs are detected in angiosperms (light grey). GH1-HMGA3 is present in both monocotyledons and dicotyledons (green), GH1-HMGA2 emerged in dicotyledons (blue), and GH1-HMGA1 is restricted to *Brassicaceae* (pink). **(B)** Schematic representation of H1 and GH1-HMGA proteins, showing the GH1 domain (grey), AT-hook motifs (orange), and the A domain (pink). **(C)** Thermodynamic analysis (ΔH°) of GH1-HMGA proteins (ParSe 2.0; ^9^), with H1.1 and H1.2 as positive controls. **(D)** Immunostaining of nuclei from 7-days old seedlings using GH1-HMGA1 and GH1-HMGA2 antibodies in WT, *gh1-hmga1*, *gh1-hmga2* and *gh1-hmga3* mutants. The signals from both antibodies (red, central panels) are distributed throughout the nucleoplasm but are excluded from the nucleolus and chromocenters. DNA was counterstained with DAPI (blue, left panels). Merged images of antibody signal and DAPI are shown on the right. Scale bar: 2 µm. **(E)** Density of peaks of GH1-HMGA1 (a - magenta), GH1-HMGA2-Myc (b - blue), GH1-HMGA3-mCherry (c - green) and TE enriched by H3K9me2 (d - black), mapped to the *Arabidopsis thaliana* TAIR10 reference genome. **(F)** Euler diagrams showing the overlap among target regions of GH1-HMGA1 (magenta), GH1-HMGA2 (blue) and GH1-HMGA3 (green).

### GH1-HMGA proteins independently bind overlapping euchromatin targets

To evaluate the functional divergence among GH1-HMGA family members and test whether they are enriched at heterochromatin like H1, we first examined their nuclear distribution in Arabidopsis seedlings using custom-made GH1-HMGA1- and GH1-HMGA2-specific antibodies (**Supplementary Figure 1C-D-E**). Immunolabeling of isolated nuclei revealed that GH1-HMGA1 and GH1-HMGA2 are distributed throughout euchromatin but are excluded from nucleoli and chromocenters (**Figure 1D**). This pattern persisted in single mutants lacking other members of the GH1-HMGA family, indicating that these proteins do not depend on each other for chromatin association. Moreover, unlike TRB proteins, another class of GH1-domain proteins that engage in heterodimeric protein-protein interactions through their coiled-coil domains ^13^, no direct interactions were detected among GH1-HMGA family members by yeast two-hybrid (Y2H) assay (**Supplementary Figure 1F**) or by co-immunoprecipitation (co-IP) (**Supplementary Figure 1G**).

To determine and compare the genome-wide binding sites of each GH1-HMGA protein, we performed ChIP-seq using the GH1-HMGA1 antibody in 7-day-old seedlings and re-analyzed public datasets for GH1-HMGA2-Myc ^8^ and GH1-HMGA3-mCherry ^12^ performed at a similar developmental stage. Peak density profiles indicate that all three GH1-HMGAs are broadly distributed across chromosome arms but absent from centromeric and pericentromeric regions, consistent with the cytological exclusion from chromocenters **(Figure 1E, Supplementary Figure 1H).** Using the same bioinformatics pipeline for the three data sets, we identified 24,577, 12,984 and 10,290 peaks, respectively, for GH1-HMGA1, −2 and −3 **(Supplementary Figure 1I-J)**. Genomic intersection analyses show that GH1-HMGA3 peaks almost entirely overlap with those of GH1-HMGA1 and GH1-HMGA2. Similarly, GH1-HMGA2 largely shares its binding sites with those of GH1-HMGA1. Yet, despite being the most recently evolved member, GH1-HMGA1 displays also more than 8,000 distinctive binding sites along the euchromatic chromosome arms (**Figure 1F, Supplementary Figure 1K**). Together, these data show that the full GH1-HMGA complement tends to cover similar genomic regions and is excluded from pericentromeric heterochromatin, which is typically occupied by H1.

### GH1-HMGAs are enriched at euchromatic TEs and play only a limited role in gene transcriptional regulation

A targeted analysis of GH1-HMGA binding at gene loci identified 3,374 genes co-enriched for GH1-HMGA1, GH1-HMGA2, and GH1-HMGA3. In contrast, a larger set of 9,794 genes is co-occupied almost exclusively by GH1-HMGA1 and GH1-HMGA2. Additionally, 5,454 genes show preferential binding by GH1-HMGA1 (**Figure 2A-B**). Interestingly, GH1-HMGA proteins preferentially localize to gene promoters and 3′ ends of genes and are largely excluded from gene bodies (**Supplementary Figure 2A**). Motif analysis of GH1-HMGA1 bound genes expectedly revealed the enrichment of GH1-HMGA1 at AT-rich sequences **(Supplementary Figure 2B),** but also at additional motifs, including “GAAGAAGAAGARAR”, a sequence closely resembling the binding sites of C2C2-DOF transcription factors, which are implicated in cold, drought, and salt stress responses (ArabidopsisDAPv1; E-value = 6.73e-02; q-value = 1.19e-01; overlap = 14 bases) ^14,15^. GO enrichment analysis of genes bound by all three GH1-HMGA proteins identified only a few significant categories, mainly related to fertilization and protein localization. Genes targeted by GH1-HMGA1 and GH1-HMGA2, but not GH1-HMGA3, showed weak enrichment for cell wall remodelling, hormone signalling, and leaf development, whereas GH1-HMGA1-specific targets showed slight enrichment for sulfur compound binding (**Supplementary Figure 2C-D**). We tested whether local enrichment of GH1-HMGA proteins impact gene expression using available RNA-seq data from *gh1-hmga12* double-mutant plants ^8^. We found no significant overlap between the set of differentially expressed and GH1-HMGA1/2-specific target genes (**Figure 2C**). Moreover, GH1-HMGA target or non-target genes had similar expression change distributions in WT and *gh1-hmga12* plants (**Figure 2D, Supplementary Figure 2E**), indicating a limited effect of GH1-HMGA1/2 dual depletion on gene expression.

**Figure 2:**
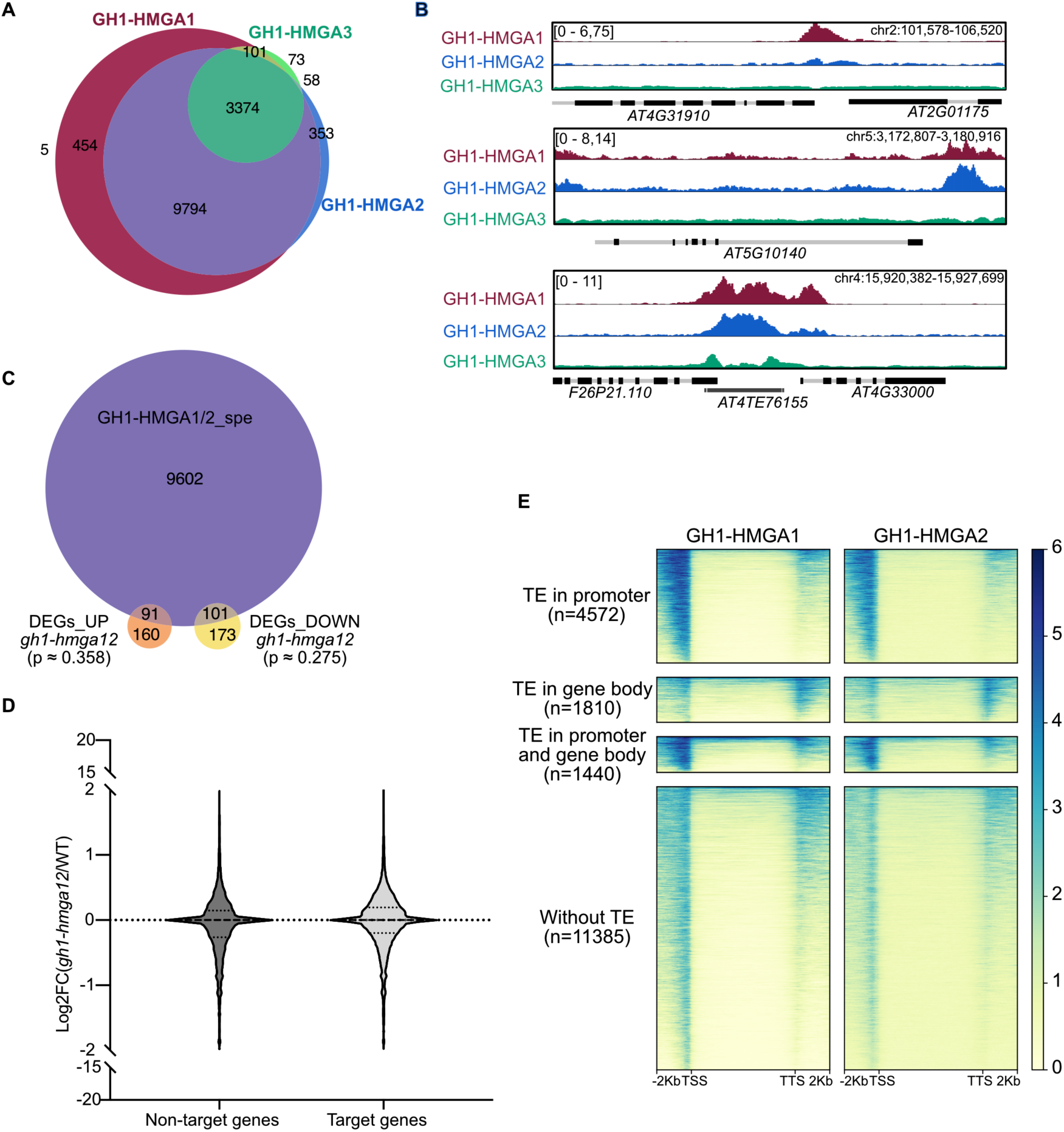
GH1-HMGA proteins preferentially bind TEs in gene-rich regions. **(A)** Euler diagram showing the overlap among target genes of GH1-HMGA1 (magenta), GH1-HMGA2 (blue) and GH1-HMGA3 (green). **(B)** Genome browser views of representative genes targeted exclusively by GH1-HMGA1 (top), jointly by GH1-HMGA1 and GH1-HMGA2 (middle), or by all three GH1-HMGAs (bottom). **(C)** Euler diagram illustrating the overlap between common target genes of GH1-HMGA1 and GH1-HMGA2 and genes up- or downregulated in *gh1-hmga1 gh1-hmga2* double mutant ^8^, p-values were calculated with a one-sided hypergeometric test. **(D)** Violin plot of mRNA log2 fold changes between WT and *gh1-hmga12* of GH1-HMGA-marked genes (n=19207) and non-target genes (n=14105) ^8^. Five Mann Whitney tests, carried out on 500 randomly selected genes of each group, were conducted, and none showed any significant difference between the target and non-target genes (**Supplementary Figure 2E**). **(E)** ChIP-seq signal for GH1-HMGA1 and GH1-HMGA2-Myc at target genes, grouped by TE presence and position.

Approximately 40% of GH1-HMGA1/2 target genes contained TEs in their promoters or gene bodies (**Figure 2E, Supplementary Figure 2F-G-H**) and genes bound by GH1-HMGA1 were significantly closer to TEs than non-targeted genes (**Supplementary Figure 2I**), suggesting that GH1-HMGA proteins preferentially associate with euchromatic TEs.

### GH1-HMGA1 preferentially binds euchromatic DNA transposons with moderate H3K9me2 and DNA methylation

This enrichment of GH1-HMGA binding at TE-containing loci led us to investigate whether these proteins directly target transposable elements more directly. A first group of TEs was found to be targeted by all three GH1-HMGA proteins, whereas a larger subset was associated only with GH1-HMGA1 and GH1-HMGA2. Unlike genes, a minor subset of TEs (n=1,517) is targeted by GH1-HMGA1 alone **(Figure 3A)**. GH1-HMGA proteins cover the entire body of target TEs **(Figure 3B-C),** the majority of which correspond to long AT-rich DNA transposons **(Figure 3D-E, Supplementary Figure 3A).** Consistent with the subnuclear distribution of GH1-HMGA1/2 proteins (**Figure 1D-E**), GH1-HMGA-targeted transposable elements displayed a strong positional bias, being preferentially located within gene-rich chromosomal regions and largely excluded from centromeres regardless of TE class **(Supplementary Figure 3B)**. In line with this distribution, GH1-HMGA proteins preferentially bound transposons with intermediate H3K9me2 levels, rather than the highly H3K9me2-enriched transposons that typify centromeric and pericentromeric heterochromatin ^16^ **(Figure 3F, Supplementary Figure 3C)**. Consistently, GH1-HMGA1-bound TEs display lower DNA methylation levels than other TEs **(Figure 3G)**. Taken together, these data show that GH1-HMGA1 preferentially binds to euchromatic DNA transposons with moderate H3K9me2 and DNA methylation levels.

**Figure 3:**
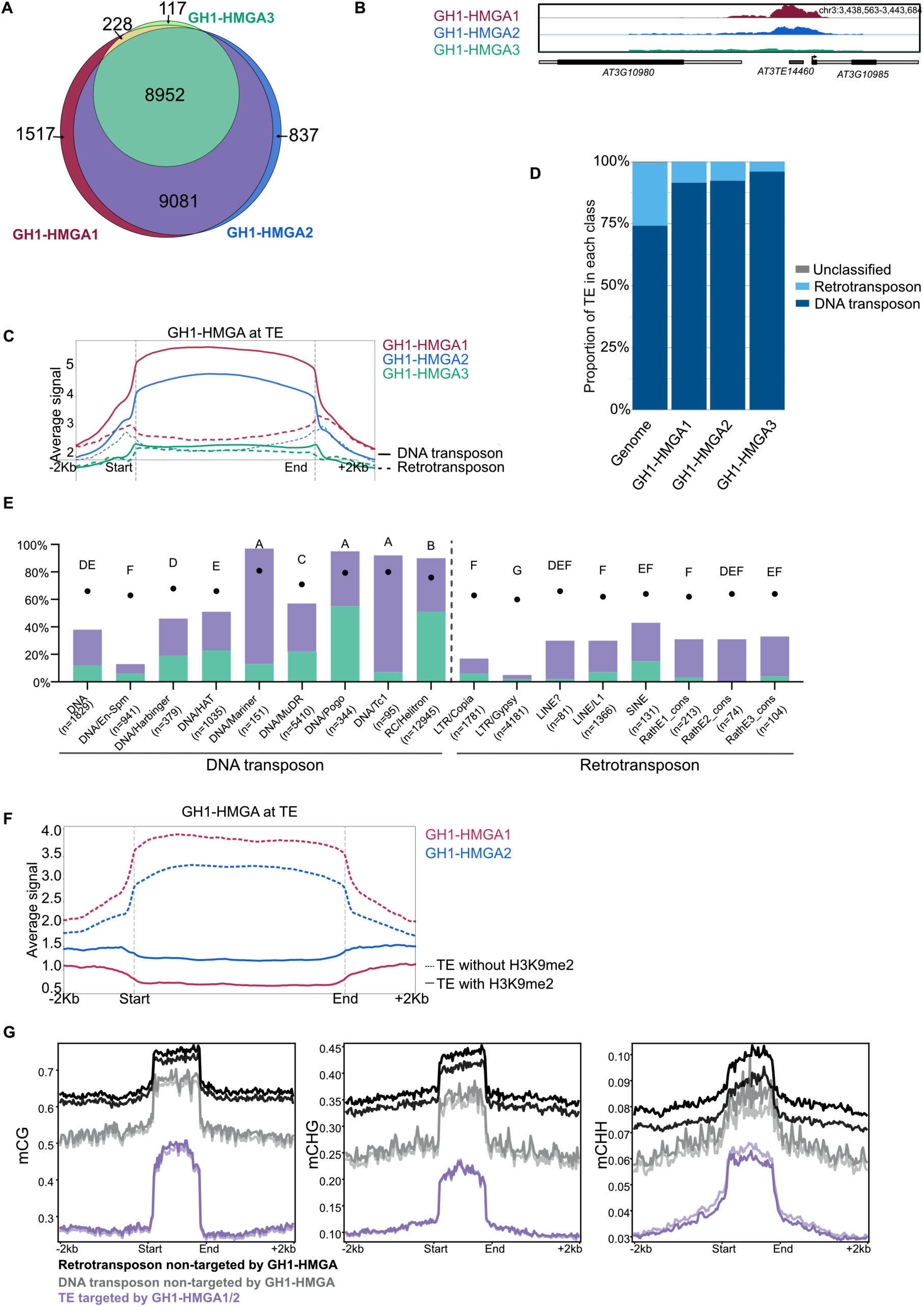
GH1-HMGA proteins preferentially bind euchromatic TEs. **(A)** Euler diagram showing the overlap among target TEs of GH1-HMGA1 (magenta), GH1-HMGA2 (blue) and GH1-HMGA3 (green). **(B)** Genome browser views of a representative TE targeted by GH1-HMGA1 and GH1-HMGA2. **(C)** Metagene plot showing ChIP-seq signal profile for each GH1-HMGA protein over its own set of target TEs. **(D)** Proportion of TE classes (retrotransposons, DNA transposons, and unclassified TEs) associated with each GH1-HMGA compared with genome-wide TE composition. Pearson’s Chi-squared tests revealed significant enrichment biases in TE class representation relative to genomic frequencies. (GH1-HMGA1: χ^2^ = 8998.6, df = 2, p < 2.2 × 10⁻¹⁶; GH1-HMGA2: χ^2^ = 8561.3, df = 2, p < 2.2 × 10⁻¹⁶; GH1-HMGA3: χ^2^ = 5180.2, df = 2, p < 2.2 × 10⁻¹⁶) (**E)** Proportion of genomic TEs targeted by GH1-HMGA1/2 or GH1-HMGA1/2/3, grouped by TE family. Bars indicate the percentage of targeted TEs within each family; green bars represent TEs targeted by GH1-HMGA1/2/3, and purple bars represent TEs targeted by GH1-HMGA1/2 only. Black dots show the median A/T content of each TE subfamily. Significant differences in A/T composition among TE families indicated in capital letters were assessed using a Dunn’s multiple comparisons test (adjusted p-value < 0.05). **(F)** Metaprofile showing ChIP-seq signal profile for GH1-HMGA1 and GH1-HMGA2 proteins at TEs enriched or not by H3K9me2. **(G)** Metaprofile showing Bisulfite-seq signal profile (mCG, mCHG, mCHH) across different groups of TEs targeted or not by GH1-HMGAs in WT plants. Two biological replicates are displayed in the same colour with different intensity.

### H1 and GH1-HMGA proteins competitively bind to TEs

Considering that H1 is enriched at heterochromatic TEs ^11^, we sought to directly compare its subnuclear and genome-wide distribution with those of GH1-HMGA proteins. Examination of chromatin occupancy of the major canonical linker histone H1.2 ^17^ revealed a marked depletion at regions targeted by GH1-HMGA1, which include not only TEs but also protein-coding genes **(Figure 4A-B-C, Supplementary Figure 4A-B-C).** Co-IP and Y2H analyses detected no interactions between the two constitutively expressed H1 variants H1.1 or H1.2 and the three GH1-HMGAs **(Supplementary Figure 1G-4D)**, altogether indicating that these two GH1-containing protein families associate with distinct target loci.

**Figure 4:**
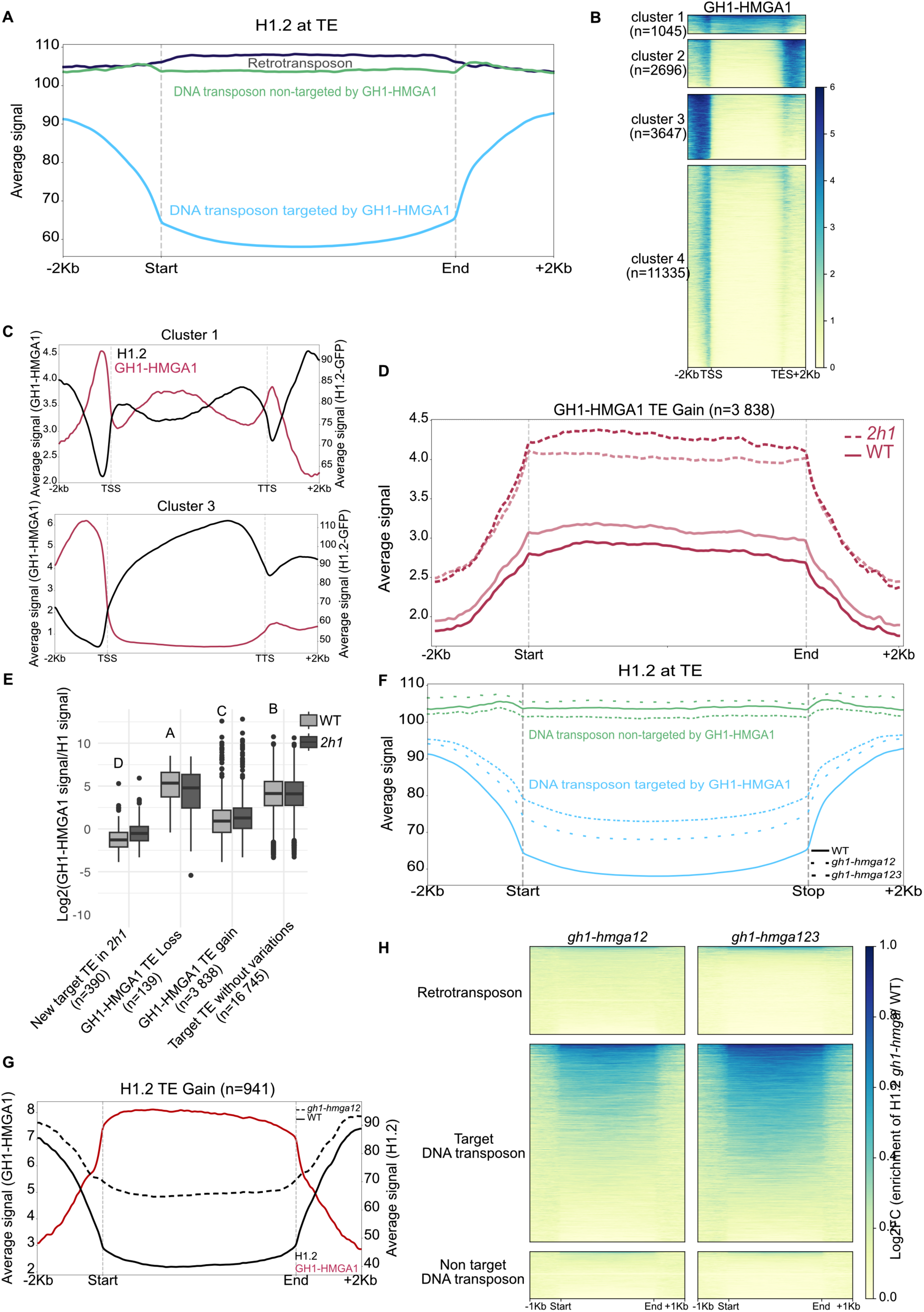
H1 and GH1-HMGA proteins exhibit distinct genomic distributions and reciprocally antagonize chromatin occupancy at euchromatic transposable elements. **(A)** Metaprofile showing H1.2-GFP ChIP-seq signal ^17^ over GH1-HMGA1 target DNA transposons (n=18149) or not targeted (n=4980) and retrotransposons (n=7931). (**B**) Heatmaps showing ChIP-seq signals of GH1-HMGA1 in WT plants at genes common to GH1-HMGA1 and GH1-HMGA2 proteins (n=9,794), following k-means clustering. **(C)** Metagene plots showing GH1-HMGA1 and H1.2 ChIP-seq enrichment for genes clusters 1 and 3 from Figure 4B. (**D)** Metaprofile showing GH1-HMGA1 ChIP-seq signal in wild-type (WT) and *2h1* mutant backgrounds over transposable elements displaying significant differential enrichment in GH1-HMGA1 as determined by DESeq2 (adjusted p-value < 0.05). Two biological replicates are displayed with different intensity. (**E**) Comparison of GH1-HMGA1 versus H1.2-GFP ChIP-seq signal log2-ratios for different transposable element groups. Boxplots show the median, interquartile range (Q1–Q3), and whiskers representing data spread excluding outliers. Significant differences in GH1-HMGA1 WT enrichments among gene and TE groups indicated in capital letters were assessed using Dunn’s multiple comparisons tests with Benjamini–Hochberg correction. **(F)** Metagene plot showing H1.2 ChIP-seq signal over TEs in *gh1-hmga12, gh1-hmga123* and WT over DNA transposon targeted or not by GH1-HMGA1. **(G)** Metaprofile showing GH1-HMGA1 ChIP-seq and H1.2 average ChIP-seq signals at TEs displaying increased H1 enrichment *gh1-hmga12* (n=941) mutants together with GH1-HMGA1-enrichment of the same targets in WT plants. **(H)** Heatmap profile showing H1 differential enrichment over retrotransposons and DNA transposons targeted or not by GH1-HMGA1 in *gh1-hmga12* or *gh1-hmga123* plants compared to WT.

To test whether H1 and GH1-HMGA chromatin enrichment is antagonistic, we probed GH1-HMGA1 distribution in an H1-deficient background by immunolabeling of isolated nuclei and ChIP-seq profiling. GH1-HMGA1 and GH1-HMGA2 immunofluorescence signals in *2h1* nuclei showed no visible distribution changes compared to wild-type nuclei **(Supplementary Figure 4E)**. ChIP-seq profiling of GH1-HMGA1 in *h1.1 h1.2* (*2h1)* plants identified an ectopic gain of GH1-HMGA1 protein at 191 genes and 390 TEs, indicating a limited but significant influence of H1 on GH1-HMGA1 genomic distribution **(Supplementary Figure 4F).** Notably, 74% of these new genes targeted by GH1-HMGA1 contain intragenic TEs, predominantly DNA transposons (78%), suggesting enhanced access of GH1-HMGA1 to euchromatic TEs upon H1 loss **(Supplementary Figure 4G-H)**. Most strikingly, in comparison to WT, GH1-HMGA1 occupancy is markedly enriched at many of its proper target genes (n=1491, 8% increased level) and TEs (n=3838, 20% increased level) in *2h1* plants **(Figure 4D, Supplementary Figure 4I-J)**, especially at loci displaying low GH1-HMGA1/H1 ratios in wild-type plants (**Figure 4E, Supplementary Figure 4K).** The expansion of the GH1-HMGA target repertoire and its increased enrichment at target genes and TEs both indicate that H1 restricts GH1-HMGA1 accumulation at multiple genes and TEs.

To explore if a competitive binding dynamic occurs between GH1-HMGA and H1, we tested whether GH1-HMGA proteins depletion promotes H1 redistribution at GH1-HMGA-associated regions. H1 immunolabeling in *gh1-hmga12 and gh1-hmga123* nuclei showed no detectable redistribution of the fluorescence signal **(Supplementary Figure 4L)**. The H1 ChIP-seq analyses in *gh1-hmga12* and *gh1-hmga123* mutants revealed that GH1-HMGA-targeted euchromatic TEs specifically gained H1 (**Figure 4F-G-H, Supplementary Figure 4M-N**). Collectively, these data support a model of reciprocal antagonism between H1 and GH1-HMGA proteins at euchromatic TEs, wherein the loss of one factor results in ectopic enrichment of the other.

### H1/GH1-HMGA interplay modulates DNA methylation landscapes at euchromatic transposable elements

Given the established role of H1 in modulating the accessibility of heterochromatic TEs to methyltransferases ^5^, we tested whether GH1-HMGA proteins similarly influence the methylation of euchromatic TEs by carrying out bisulfite sequencing (BS-seq) of genomic DNA extracted from *gh1-hmga12* and *gh1-hmga123* mutant seedlings. Consistent with the absence of GH1-HMGA proteins at heterochromatic TEs, DNA methylation levels remained largely unaffected at these loci in both mutant lines (**Figure 5A**). In contrast, in both mutants we observed a robust increase in CHG and CHH methylation at euchromatic TEs (**Figure 5A**), particularly those bound by all three GH1-HMGAs (**Figure 5B**, **Supplementary Figure 5A-B**). Notably, combined loss of GH1-HMGA1 and GH1-HMGA2, irrespective of GH1-HMGA3 binding, led to a comparable CHG and CHH hypermethylation at shared targets, indicating that GH1-HMGA1 and GH1-HMGA2 are the primary factors restricting non-CG methylation at these loci.

**Figure 5:**
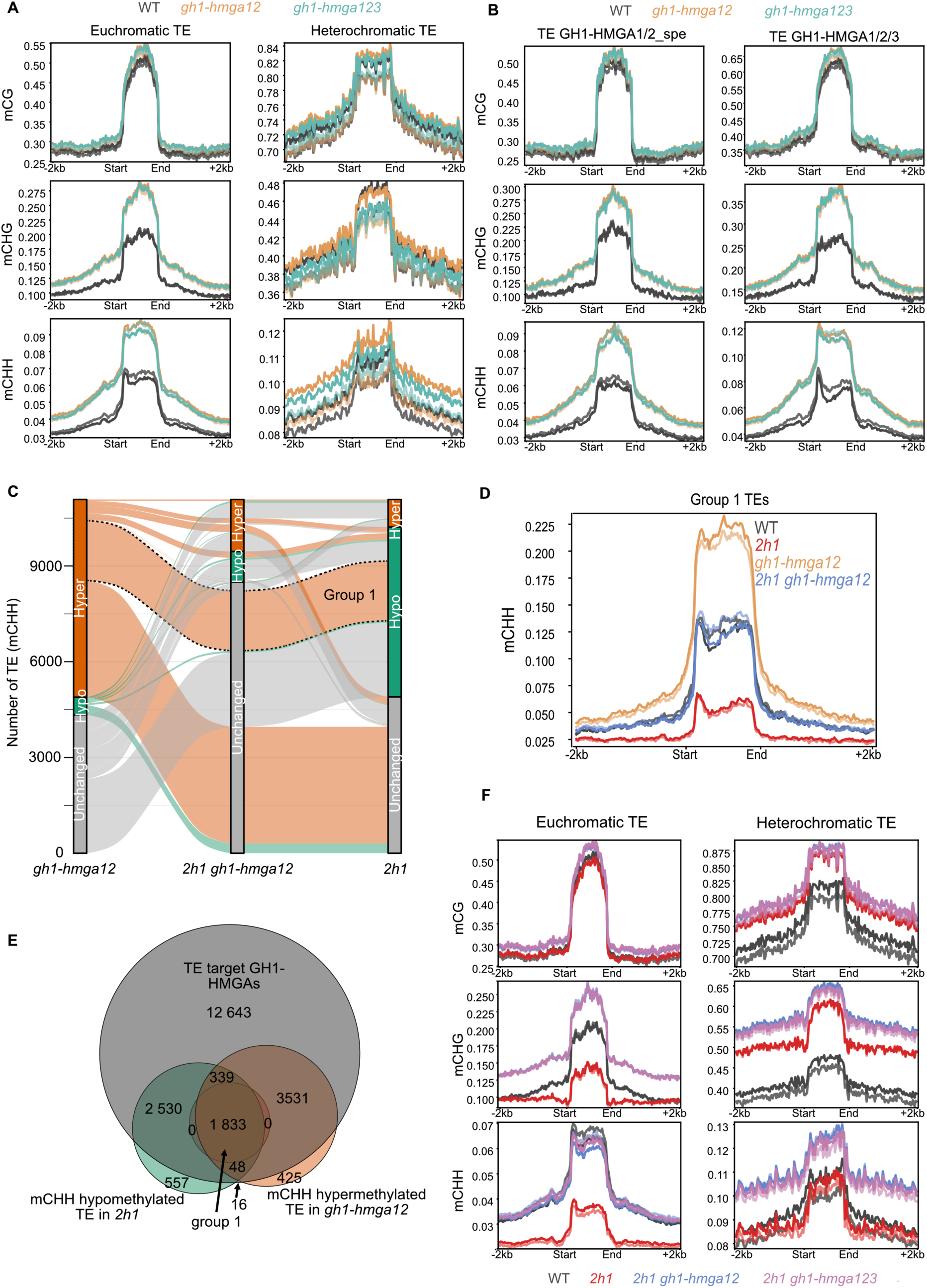
Antagonistic interplay between H1 and GH1-HMGA proteins controls DNA methylation at euchromatic transposons. **(A)** Metaprofile showing Bisulfite-seq signal profile (mCG, mCHG, mCHH) across euchromatic and heterochromatic transposable elements for WT, *gh1-hmga12* and *gh1-hmga123*. Biological replicates are displayed in the same colour with different intensity. **(B)** Metaprofile showing Bisulfite-seq signal profile (mCG, mCHG, mCHH) across TE targets shared between GH1-HMGA1 and GH1-HMGA2, or common to all three GH1-HMGAs for WT, *gh1-hmga12* and *gh1-hmga123*. Biological replicates are displayed in the same colour with different intensity. (**C)** Alluvial diagram of CHH methylation changes at TEs in *2h1, gh1-hmga12*, and *2h1 gh1-hmga12* mutants. Hypermethylated, hypomethylated, and unchanged TEs relative to WT are shown in orange, green, and grey, respectively. TEs hypermethylated in *gh1-hmga12*, hypomethylated in *2h1*, and unchanged *in 2h1 gh1-hmga12* relative to WT are outlined with a dotted line. **(D)** Metaprofile of Bisulfite-seq mCHH signal across the TE group 1 in (C). Two iological replicates are displayed in the same color with different intensity. **(E)** Euler diagram showing overlap between GH1-HMGA target TEs and the TEs either CHH hypomethylated in *2h1* or CHH hypermethylated in *gh1-hmga12*. **(F)** Metaprofile showing Bisulfite-seq signal profile (mCG, mCHG, mCHH) across euchromatic and heterochromatic transposable elements for WT, *2h1, 2h1 gh1-hmga12* and *2h1 gh1-hmga123*. Two biological replicates are displayed in the same colour with different intensity.

To investigate the impact of antagonistic binding between GH1-HMGA and H1 proteins on DNA methylation, we compared DNA methylation changes induced by H1 or GH1-HMGA depletion alone or in combination. Consistent with previous reports ^4,5^, the *2h1* mutant showed ∼5,000 hypomethylated TEs compared to WT plants (**Figure 5C, Supplementary Figure 5C**). By contrast, *gh1-hmga* mutants mainly showed CHG and CHH hypermethylation compared to WT, affecting about 5,500 TEs. Importantly, most of this hypermethylation was suppressed in plants combining H1 and GH1-HMGA loss-of-function mutations. This suppressive effect was also observed for a subset of about 2,000 TEs that are direct GH1-HMGA targets (Group 1) and which are hypermethylated in *gh1-hmga* mutants, but hypomethylated in *2h1* **(Figure 5D-E - Supplementary Figure 5D-G)**. Conversely, CHG and CHH hypomethylation at euchromatic TEs caused by H1 loss was fully rescued in the *2h1 gh1-hmga12* and the *2h1 gh1-hmga123* mutant plants, CHG methylation further exceeding that of wild-type plants at some TEs **(Figure 5F)**. By contrast, H1-dependent hypermethylation persisted at heterochromatic TEs in the *2h1 gh1-hmga12* mutants, which is consistent with the absence of GH1-HMGA chromatin association at this set of TEs. Overall, these results indicate that GH1-HMGA proteins act in concert with H1 to dampen DNA methylation at euchromatic TEs.

### GH1-HMGA1 and GH1-HMGA2 restrict RdDM-mediated DNA methylation at H1-poor euchromatic TEs

Given that GH1-HMGA proteins are primarily associated with euchromatic TEs carrying moderate H3K9me2 levels, and that Differentially Methylated Regions (DMRs) detected in GH1-HMGA mutants are more enriched in RdDM than in CMT2 targets (**Supplementary Figure 6A**), their CHG and CHH methylation may reflect RdDM activity ^1^. To test whether GH1-HMGA proteins affect RdDM activity, we profiled small RNAs in *gh1-hmga12* seedlings. The sRNA-seq analyses showed that small RNAs are globally enriched at *gh1-hmga12* hypermethylated loci (**Figure 6A-B**), suggesting that GH1-HMGA12 restrict sRNA production of associated TEs, for example, by hindering their DNA accessibility. To functionally test whether the RdDM pathway underlies the hypermethylation observed at euchromatic TEs upon GH1-HMGA12 loss, we generated a *nrpe1 gh1-hmga12* triple mutant line in which *NRPE1* gene, encoding the RNA Polymerase V subunit essential for RdDM, is disrupted. BS-seq analysis of DNA from WT and *nrpe1 gh1-hmga12* mutant seedlings revealed that RdDM is essential for a large part of TE hypermethylation observed upon GH1-HMGA12 loss, as its level is strongly reduced in the absence in *gh1-hmga12* plants (**Figure 6B-C**). Collectively, these results show that GH1-HMGA1 and GH1-HMGA2 are enriched at euchromatic TEs depleted in H1, where they modulate DNA methylation by restricting RdDM, limiting small RNA accumulation and preventing excessive CHG and CHH methylation.

**Figure 6:**
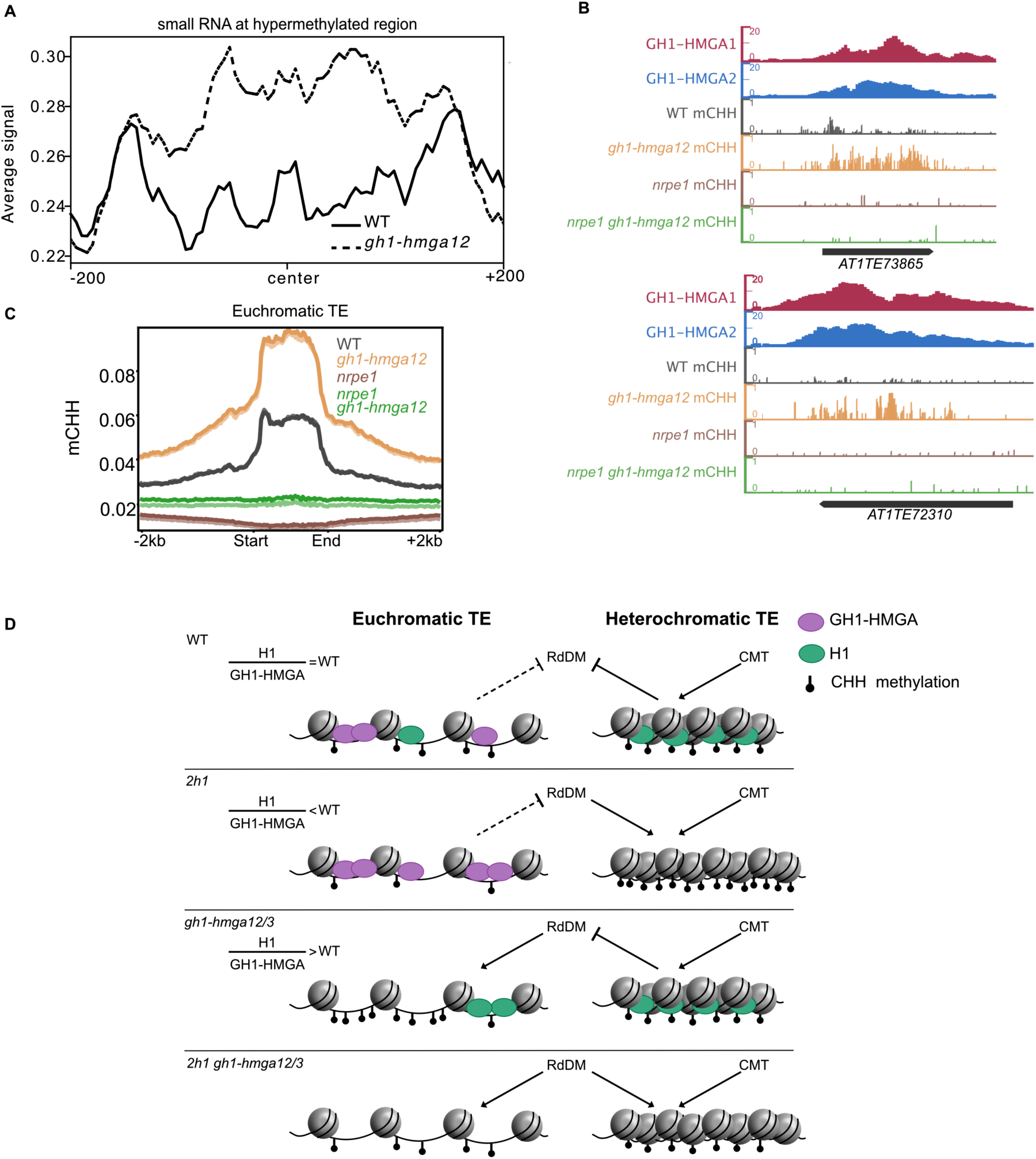
GH1-HMGA proteins restrict RdDM-mediated DNA methylation at H1-poor DNA transposons. **(A)** Average small RNA signal centered on hypermethylated DMRs at TE targets of GH1-HMGA. **(B)** Genome browser view at transposons exhibiting mCHH hypermethylation in the *gh1-hmga12* mutant and targeted by GH1-HMGA1, shown for wild type (WT), *gh1-hmga12, nrpe1* and *nrpe1 gh1-hmga12* mutants. **(C)** Metaprofile showing Bisulfite-seq signal profile of mCHH at euchromatic TEs for WT, *gh1-hmga12, nrpe1* and *nrpe1 gh1-hmga12.* Two biological replicates are displayed in the same color with different intensity. **(D)** Working model of H1/HMGA-dependent regulation of TE DNA methylation. We propose that the balance between H1 (green) and GH1-HMGA (purple) proteins partitions euchromatic and heterochromatic TE domains and thereby differentially modulates the access of RdDM- and CMT-dependent methylation pathways. In wild-type plants, GH1-HMGA1 is enriched at euchromatic TEs, whereas H1 accumulates at heterochromatic TEs. In *2h1* and *gh1*-*hmga12 or gh1*-*hmga123* backgrounds, perturbation of this balance alters GH1-HMGA1/H1 distribution and is associated with changes in TE methylation. In the *2h1 gh1-hmga12/3* mutant, the combined loss of H1 and GH1-HMGA1 relieves RdDM from these constraints and results in a broad redistribution of CHH methylation. Arrows indicate the proposed effects of H1/GH1-HMGA1 balance on RdDM- and CMT-dependent methylation pathways in each chromatin context.

## Discussion

Here, we show that GH1-HMGA and linker histone H1 homeostasis define a complementary chromatin regulatory system that helps partition DNA methylation pathways. In this framework, H1 primarily constraining CMT2/CMT3-associated silencing in heterochromatin while GH1-HMGAs modulate RdDM at euchromatic TEs. This model explains the hypomethylation phenotype observed at euchromatic TEs in *2h1* mutants, which may reflect altered balance between H1 and GH1-HMGAs at shared chromatin targets rather than a simple reduction in RdDM activity. Here, H1 acts as the dominant barrier in heterochromatin, whereas GH1-HMGAs restrict RdDM in euchromatic regions, where they preferentially associate with AT-rich linker DNA. Removing both barriers, as in plants lacking H1 and GH1-HMGA1 release RdDM constraints and broaden CHH methylation across the genome **(Figure 6D)**. Together, these observations support a model in which H1 and GH1-HMGAs act as partially complementary players that help maintain epigenome integrity by limiting aberrant RdDM activity.

The extensive overlap between regions targeted by all three GH1-HMGA proteins occurs without detectable protein-protein interactions, suggesting that they can independently recognize shared genomic regions. However, this apparent co-occupancy could also reflect cell-to-cell or temporal exclusivity that is not captured by our experiments. Although H1 lacks a sequence-specific DNA binding domain, the AT-hook motifs of GH1-HMGA proteins are consistent with their preference for A/T-rich sequences ^8,18^. This suggests that local sequence composition contributes to their genomic distribution, although TE size and chromatin context may also influence occupancy. Additionally, the rapid residence times and dynamic chromatin scanning, a well-characterized behaviour of mammalian GH1-HMGA proteins, may similarly allow GH1-HMGA to outcompete H1 for binding to AT-rich linker DNA ^19–22^. With an additional GH1 DNA-binding domain, GH1-HMGA proteins may stably anchor at linker DNA, contributing to antagonistic chromatin association with H1. Consistent with observations for GH1-containing TRB proteins ^17,23^ our data support a model in which GH1-HMGA proteins compete with H1 for GH1-mediated chromatin association, thereby limiting H1 enrichment over euchromatic TEs.

Importantly, this antagonistic chromatin occupancy has direct functional consequences for RdDM activity at euchromatic TEs. Loss of GH1-HMGA1 and GH1-HMGA2 leads to enhanced small RNA accumulation together with increased CHG and CHH methylation at euchromatic TEs. Critically, this hypermethylation is strongly dependent on NRPE1, indicating that GH1-HMGA proteins restrict RdDM at these loci. By contrast, heterochromatic TEs remain largely unaffected by GH1-HMGA depletion, consistent with the absence of GH1-HMGA enrichment in these regions and the dominant role of the CMT2/CMT3 pathway for their silencing. These findings support a model in which GH1-HMGA proteins limit access of the RdDM machinery to euchromatic TE chromatin, potentially by reducing recruitment or activity of RNA polymerase IV and its associated SNF2-like CLASSY (CLSY) factors particularly CLSY1/2. This mechanism aligns with recent findings on REPRESSOR OF SILENCING 1 (ROS1), where occupancy-based mechanisms maintain hypomethylation largely independently of catalytic activity ^24^. Thus, GH1-HMGAs appear to modulate the extent of DNA methylation rather than establishing extreme methylation states.

Several additional features of GH1-HMGA proteins may further explain how they antagonize RdDM at these elements. For instance, thermodynamic profiling revealed that GH1-HMGA1 and GH1-HMGA2 are preditected have a high propensity for LLPS, comparable to that of H1, whereas GH1-HMGA3 does not. This difference is consistent with the observation that GH1-HMGA1 and GH1-HMGA2, but not GH1-HMGA3, make the major contribution to RdDM antagonism at euchromatic TEs. One possibility is that local condensate-forming properties help GH1-HMGA1/2 to stabilize a chromatin environment that is permissive to their own occupancy but less accessible to RdDM factors. Their evolutionary trajectory suggests euchromatic specialization of GH1-HMGA1/2, possibly adapting to the integration of multiple TEs in gene-rich regions ^5,25^. Whether LLPS directly drives TE-specific RdDM antagonism remains to be functionally tested.

Beyond its immediate impact on methylation, this balance between H1 and GH1-HMGA proteins may have broader implications for genome function and adaptation. RdDM is widely viewed as a surveillance pathway acting preferentially on euchromatic TEs, particularly those inserted near genes, where excessive methylation spreading could have deleterious regulatory consequences ^2526^. In this context, GH1-HMGA proteins may help prevent irreversible silencing of specific euchromatic TEs, thereby preserving their potential to influence on neighbouring gene expression. Such a function may be especially relevant under environmental stress, where transient TE activation can promote regulatory and genetic diversity. By limiting excessive CHG and CHH methylation while still maintaining TE control, GH1-HMGA proteins may therefore contribute to a balance between epigenome stability and regulatory plasticity. This possibility is particularly intriguing given the evolutionary emergence of GH1-HMGA1 and GH1-HMGA2 in angiosperms and dicots, suggesting that these proteins may have specialized to fine-tune TE control in gene-rich chromosomal environments.

## Methods

### Plant materials and growth conditions

Seeds were surface-sterilized using 70% ethanol containing 0.01% SDS, plated on agar-solidified Murashige and Skoog (MS) medium supplemented with 0.9% agar and 1% sucrose, and stratified in the dark at 4°C for 2 days. For ChIP experiments targeting GH1-HMGA1 or H1, seedlings were grown on full-strength (1X) MS medium. Seedlings were grown under long-day conditions (16 hours light / 8 hours dark) at 23°C in a growth chamber. Seedlings were harvested at 5 days after germination unless otherwise indicated.

The *gh1-hmga* mutants used in this study were previously described by ^8^ and were generated by crossing the *gh1-hmga1* (AT3G18035 - SALK_071403), *gh1-hmga2* (At1g48620 - SALK_116292), and *gh1-hmga3* (AT1G14900 - SALK_078336) mutant lines. The *2h1* mutants were obtained by crossing the *h1.1* (AT1G06760 - SALK_N628430; Rea, 2012) and *h1.2* (AT2G30620 - GK-116E080; Rutowicz, 2015) mutant lines. The *2h1 gh1-hmga12* and *2h1 gh1-hmga123* mutants were generated by genetic crosses. The *nrpe1 (*AT2G40030 - SALK_029919) was first described in ^27^. The *pGH1-HMGA1::GH1-HMGA1* lines used for co-immunoprecipitation experiments were previously described in ^7^.

### Phylogenetic studies

Phylogenetic trees were constructed using protein sequences from species representing the whole plant lineage: *Chlamydomonas reinhardtii, Oestreococcus tauri, Marchantia polymorpha, Physcomitrella patens, Selaginella moellendorffii, Pinus taeda, Picea sitchensis, Amborella trichopoda, Nymphaea colorata, Musa acuminata, Ananas comosus, Zea mays, Sorghum bicolor, Triticum aestivum, Oryza sativa, Solanum lycopersicum, Vitis vinifera, , Populus trichocarpa, Glycine max, Prunus persica, Theobroma cacao, Arabidopsis thaliana, Arabidopsis lyrata, Brassica rapa and Eutrema salsugineum*.

A workflow was implemented as a Bash pipeline to identify homologous sequences by reciprocal best hits using MMseqs2 (threshold 1e-10) ^28^, and then to extract the corresponding sequences from each database. The selected sequences were grouped by orthologous set, aligned with MAFFT 7.407 ^29^ using the parameters --globalpair --maxiterate 16 --auto --inputorder, and used to infer maximum-likelihood trees with IQ-TREE v2.2.0.3 ^30^ with the LG substitution model and 1000 bootstrap replicates. Trees were refined using the Interactive Tree Of Life ^31^. More details and the exact implementation are available in the workflow script: https://gitlab.com/lilian_fau/fhb_secure//blob/main/post_treatments/script_07_PHYLOGENY.sh?ref_type=heads. Protein accession numbers and sequences used in this study are listed in Supp Table 1.

### Y2H assay

Yeast cultures were grown at 30°C on YPD or on selective SD medium. Full-length coding sequences were cloned into bait (pDEST-GBKT7) or prey (pDEST-GADT7) vectors ^32^ then transformed into *Saccharomyces cerevisiae* strains AH109 Gold and Y187 (Clontech, MATCHMAKER GAL4 Two-Hybrid System), respectively, using a classical heat-shock protocol ^33^ and grown on selective medium lacking Trp or Leu. The two yeast strains were mated on YPD, and diploids were selected on SD-Leu-Trp. Protein–protein interactions were detected by growth on low-stringency selective medium lacking Leu, Trp, and His. Empty pDEST-GBKT7 or pDEST-GADT7 vectors were used as negative controls.

### Antibodies production

Rabbit polyclonal antibodies were raised against two synthetic peptides derived from GH1-HMGA1 and affinity-purified by Covalab (Lyon, France), and against one synthetic peptide derived from GH1-HMGA2. See **Supplementary Figure 1** for peptide positions and sequences, as well as antibody validation.

### Slide preparation and immunofluorescence staining

Immunostaining of GH1-HMGA1, GH1-HMGA2 and H1 was performed as described in ^34^. Antibodies and dilutions used in this study are described in **Supp_data_6**.

### Co-immunoprecipitation

Co-immunoprecipitation assays were performed on 7-day-old seedlings of *Arabidopsis* transgenic lines expressing GH1-HMGA1-GFP, essentially as previously described in ^35^, with some modifications depending on the experiment. Following extraction with Nuclear extraction buffer 1 (NEB1) buffer (0.4 M sucrose, 10 mM pH 8.0, 10 mM MgCl_2_, 5 mM BME, 0.1 mM PMSF, and one cOmplete™ Mini EDTA-free Protease Inhibitor Cocktail tablet (Roche), in some cases, samples were crosslinked with 1% formaldehyde (Thermo Scientific, 28906) in a modified NEB2 buffer (0.25 M sucrose, 60 mM HEPES pH 8.0, 10 mM MgCl_2_, 0.3% Triton X-100, 0.1 mM PMSF, and one cOmplete™ Mini EDTA-free Protease Inhibitor Cocktail tablet (Roche) for 8 minutes at room temperature. Crosslinking was quenched by the addition of glycine to a final concentration of 0.125 M, followed by centrifugation at 10,000 g for 10 minutes at 4°C. Pellets were washed several times with NEB3 buffer (1.7 M sucrose, 10 mM Tris-HCl pH 8.0, 2 mM MgCl2, 0.15% Triton X-100, 5 mM BME, 0.1 mM PMSF and one cOmplete™ Mini EDTA-free Protease Inhibitor Cocktail tablet (Roche)), and subsequently centrifuged at high speed for 1 hour at 4°C. At the end of the extraction process, nuclear lysis was performed using Nuclei Lysis Buffer (NBL) (50 mM Tris-HCl, pH 8.0, 10 mM EDTA, 0.1% SDS, 0.1 mM PMSF, one cOmplete™ Mini EDTA-free Protease Inhibitor Cocktail tablet (Roche)), with a 1-hour incubation. Depending on the experiment, samples were subsequently treated with 87.5 units of Benzonase® Nuclease (Millipore) at 4°C for 1 hour. This step was included or omitted according to specific experimental requirements. Immunoprecipitation was then carried out using ChromoTek GFP-Trap® Magnetic Agarose beads, with overnight incubation at 4°C under constant rotation. The efficiency of the GFP-based immunoprecipitation was verified using an anti-GFP antibody (Invitrogen, #A-11122). Detection of GH1-HMGA1 and GH1-HMGA2 as potential interactors was performed using specific antibodies described in (**Supplementary Figure 1**).

### RNA extraction

Total RNA was extracted from adult leaves of 3- to 4-week-old Arabidopsis plants grown in soil. For tissue disruption, flash-frozen leaf samples were ground twice in 2 ml tubes using a Tissue Lyser (Qiagen) at 30 Hz for 30 seconds, repeated twice. RNA was then isolated using the RNeasy Plant Mini Kit (Qiagen), following the manufacturer’s instructions.

### ChIP-seq

GH1-HMGA1 and H1 ChIP-seq profiling was performed essentially as described in ^17^, with slight modifications. Approximately 1 g of 7-day-old *in vitro*-grown *Arabidopsis* seedlings was crosslinked with 1% formaldehyde (Sigma-Aldrich) under vacuum for 8 minutes, repeated twice. Crosslinking was quenched with 0.125 M glycine. Fixed tissue was then ground to a fine powder in liquid nitrogen using a mortar and pestle. Nuclei were isolated and lysed in Nuclei Lysis Buffer (NBL) containing 0.1% Triton X-100. For GH1-HMGA1 ChIP, chromatin was sheared using a Covaris S220 Focused-ultrasonicator for 15 minutes (peak power 110 W, duty factor 5%, 200 cycles per burst). For H1 ChIP, chromatin was sheared using a Bioruptor Pico. Immunoprecipitation of 100 µg (protein content) chromatin pre-cleared by 20µL of uncoupled Dynabeads™ Protein A was performed either using a GH1-HMGA1-specific antibody or Agrisera anti-H1 (#AS111801) coupled to 35 µL of Dynabeads™ Protein A (Invitrogen) per sample. As control for immunoprecipitation specificity, chromatin from the *gh1-hmga1* mutant line, or the *h1.1 h1.2* (*2h1*) double mutant line lacking both canonical linker histones, were processed in parallel. Immunoprecipitated DNA was purified using Zymo ChIP DNA Clean & Concentrator columns and quantified using a Qubit fluorometer (Thermo Fisher). GH1-HMGA1 and H1 ChIP-seq libraries were prepared using the Illumina TruSeq ChIP Sample Preparation Kit or the NEBNext® Ultra™ II DNA Library Prep Kit for Illumina® (Catalog No. E7645), respectively, and sequenced on a DNBSEQ-G400 platform (BGI) as single-end 50 bp or paired-end 100bp, respectively. Each ChIP-seq experiment was performed in two biological replicates.

### Bioinformatics for ChIP-seq analysis

Raw sequencing reads were pre-processed by the sequencing platform. Reads of GH1-HMGA’s ChIP-seq ^8,12^ were mapped to the *Arabidopsis thaliana* TAIR10 genome using Bowtie2 (Galaxy version 2.5.3+galaxy1) with the --very-sensitive setting. Duplicate reads were marked using MarkDuplicates (Galaxy version 3.1.1.0). Peak calling was performed with MACS2 (Galaxy version 2.2.9.1+galaxy0) using the following settings: --nomodel, --broad, --qvalue 0.01, and an effective genome size of 1.2e8. Peaks common to both biological replicates were retained using bedtools intersect (v2.31.1). Mitochondrial (ChrM) and chloroplast (ChrC) reads were excluded from the analysis. Signal normalization was performed using S3norm on bedgraph files, and signal tracks used for visualization corresponded to S3norm-normalized negative log10(p-value) files ^36^. Downstream analyses, including annotation of peaks and signal quantification over genomic features, were performed using bedtools v2.31.1 on the command line. The comparison of GH1-HMGA protein occupancy across the TAIR10 genome sequence was performed by merging peaks obtained for each protein using bedtools merge (bedtools v2.31.1) to define regions of enrichment. Peaks from each protein were then intersected with these regions, allowing the construction of a Euler diagram representing shared and specific binding sites. H1 ChIP data were processed similarly, using partially different filtering and normalization parameters, following the pipelines described in ^37,38^. GH1-HMGA and H1 signals were normalized with S3norm ^36^. Log2 fold-change (log2FC) ratios were computed from normalized signals, and signal tracks used for visualization corresponded to S3norm-normalized negative log10(p-value) bedgraph files.

### Whole Genome Bisulfite Sequencing

DNA extraction was performed using the Promega kit (Wizard® Genomic DNA Purification Kit). For each sample, bisulfite sequencing reads were quality-trimmed and adapter-removed using fastp v0.22 with default parameters, then aligned to the TAIR10 reference assembly using Bowtie2 v2.2.5 with default parameters. Duplicate reads were removed using Picard MarkDuplicates v2.18, and per-cytosine DNA methylation was subsequently called using the Bismark methylation extractor v0.22.3 with default parameters.

### sRNA extraction and Sequencing

The precipitation of small RNA fraction was performed following RNAzol® RT method (Molecular Research Center). For each sample, small RNA sequencing reads were processed following a previously described methodology ^26^. Briefly, adapters were removed using fastp v0.22 with default parameters, and reads were aligned to the TAIR10 reference assembly using Bowtie2 v2.2.5 with the following parameters: --end-to-end --very-sensitive -L 15 -N 1. Reads mapping to nuclear chromosomes with sizes ranging from 15 to 30nt were extracted using SAMtools v1.21 and categorized into 21nt or 23-24nt size classes based on their length. The reads in the 23-24 class were converted into per-base coverage tracks using bamCoverage v3.5, with a bin size of 1bp and normalized to reads per million (RPM) using a scaling factor computed from the total number of nuclear reads with sizes ranging from 15 to 30nt.

## Funding

This work was primarily supported by grant ANR-22-CE20-0001 from Agence Nationale de la Recherche (ANR, France) to S.A, L.Q., and F.B. It was also supported by the CNRS program EPIPLANT and the COST Action INDEPTH.

## Supplementary files

Supp_data_1.xlsx : Informations related to Figure 1 and Supplementary Figure 1

Supp_data_2.xlsx : Informations related to Figure 2 and Supplementary Figure 2

Supp_data_3.xlsx : Informations related to Figure 3 and Supplementary Figure 3

Supp_data_4.xlsx : Informations related to Figure 4 and Supplementary Figure 4

Supp_data_5.xlsx : Informations related to Figure 5 and Supplementary Figure 5

Supp_data_6.xlsx : Informations related to Primers, vectors and Antibodies used in this study

## Acknowledgements

We thank the technical platforms of iGReD for their support, in particular the Bioinformatics Platform (BIM), as well as the CLIC, the Anipath histopathology platform, and the plant platforms.

---

**Supplementary figure 1:**
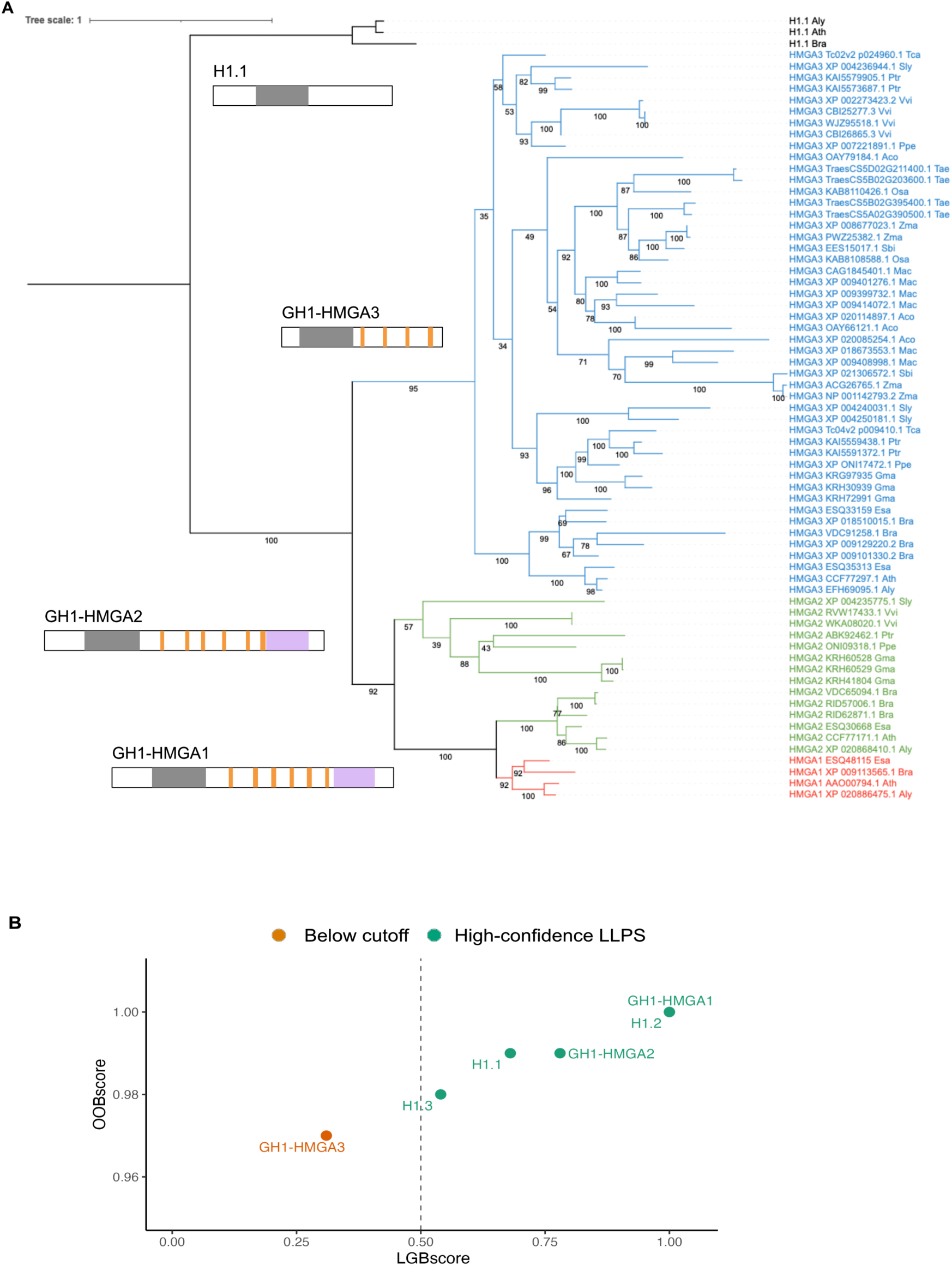
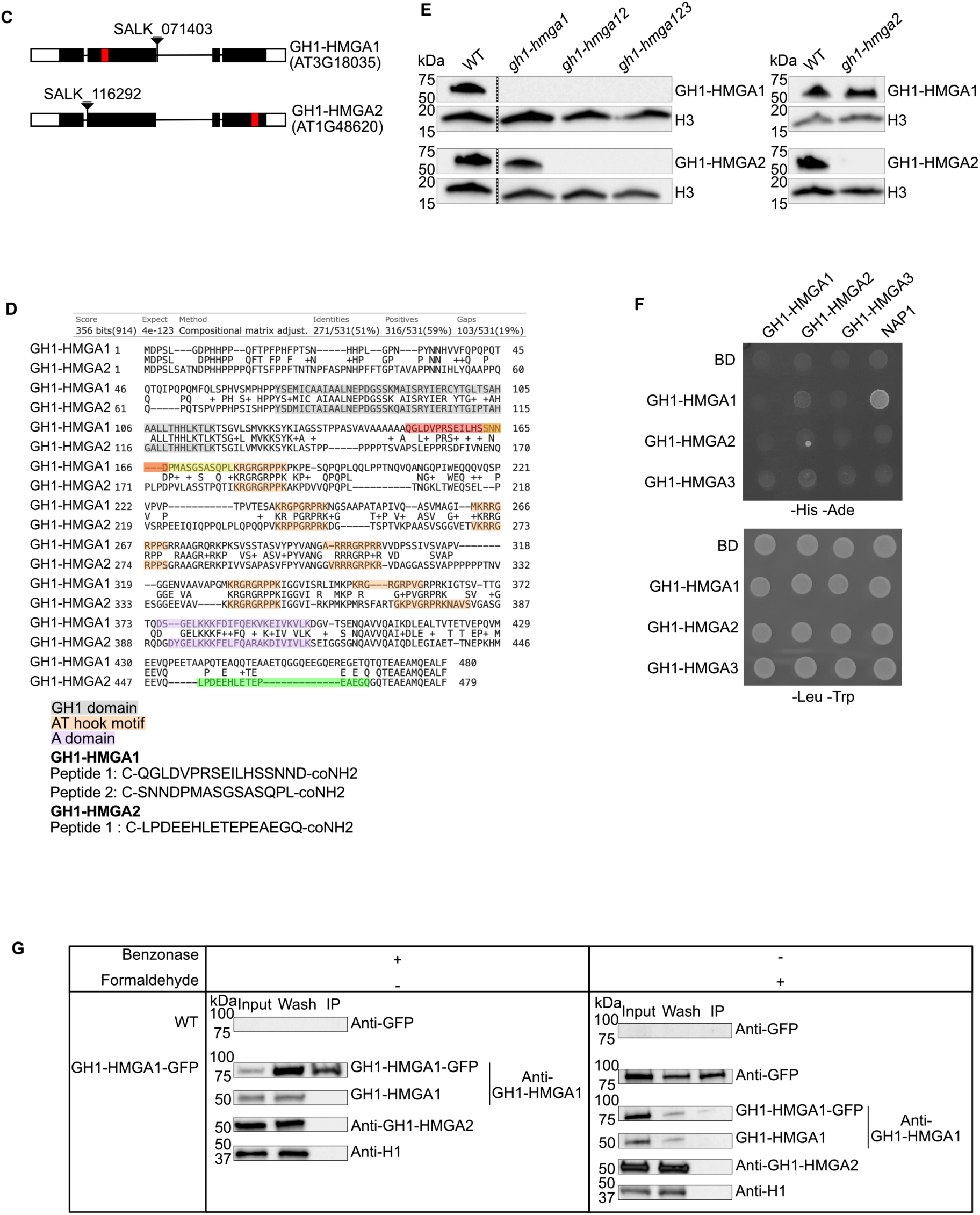
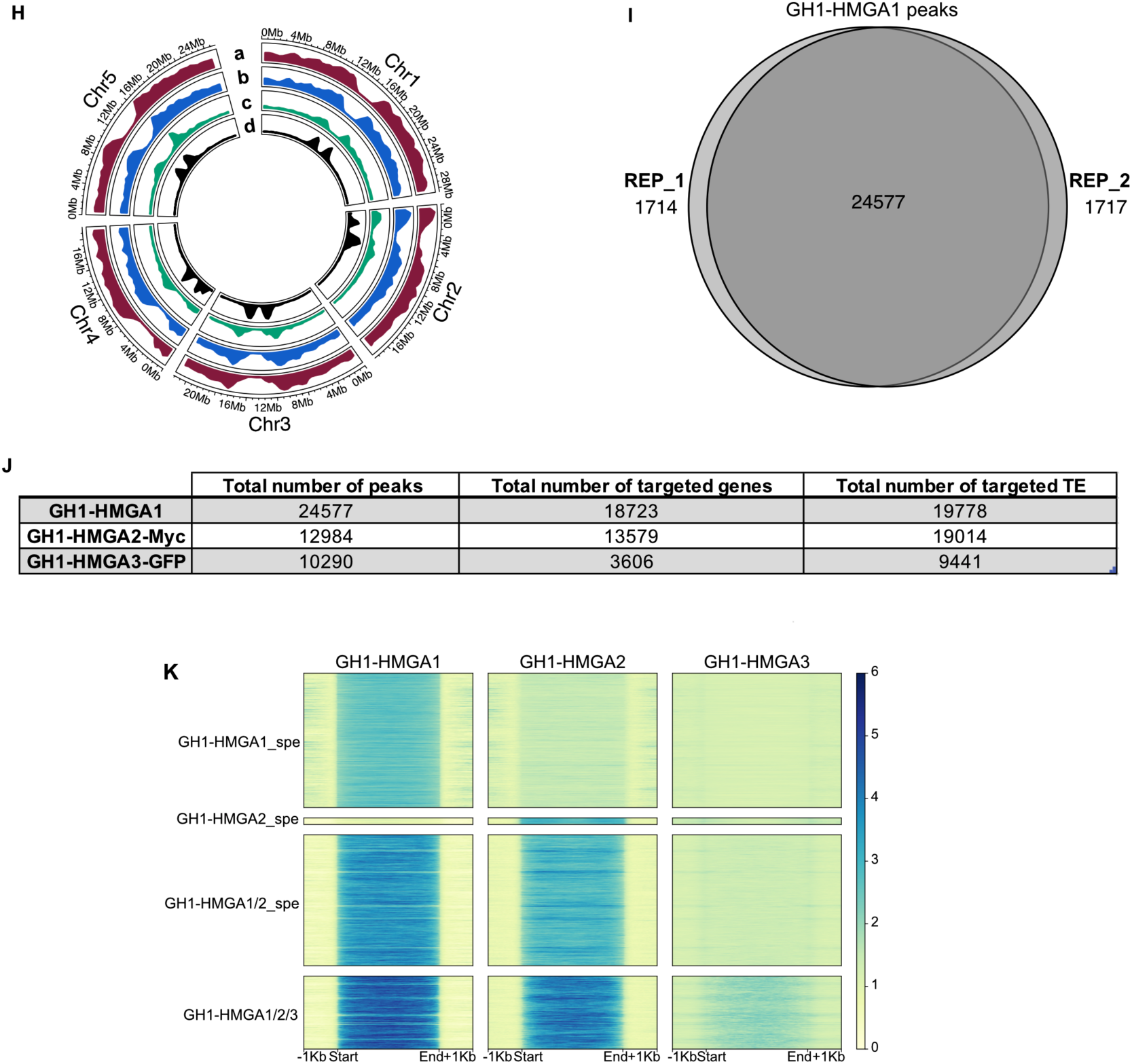
(**A**) Rooted maximum likelihood phylogenetic tree for GH1-HMGA orthologs from 25 plant species. Bootstrap values are indicated for each branch. **(B)** Scatter plot of LGBscore versus OOBscore for H1.1, H1.2, H1.3, GH1-HMGA1, GH1-HMGA2, GH1-HMGA3. Proteins with LGBscore > 0.5 and OOBscore > 0.9 were classified as high-confidence LLPS candidates, following the cutoff used in ^10^. (**C**) Gene models of GH1-HMGA1 and GH1-HMGA2 showing exon–intron structure. The positions of T-DNA insertions in the respective mutant lines are indicated, along with the regions encoding the epitopes recognized by each antibody (red). **(D)** Schematic representation of the GH1-HMGA1 and GH1-HMGA2 protein sequences, highlighting the GH1 domain (grey), AT-hook motifs (orange), and the A-domain (violet). Peptides used for antibody generation are shown in red, yellow and green. **(E)** Western blot analysis using the respective antibodies on wild-type (WT) and mutant lines of 7-days old seedlings confirms the specificity of each antibody. **(F)** Interaction among the three GH1-HMGAs and NAP1 probed in the Y2H system. Growth on selective medium lacking histidine and adenine reveals interaction between the two proteins tested (left panel) and on synthetic medium lacking leucine and tryptophan, selecting for the presence of the bait and prey vectors (right panel). Vertical, translational fusion with the Gal4-DNA-binding domain (BD). BD indicate the empty vector. NAP1 serves as positive control according to ^39^. **(G)** Co-immunoprecipitation (Co-IP) assays testing interaction between GH1-HMGA1 and GH1-HMGA2. Proteins isolated from 7-d old seedlings of pGH1-HMGA1::GH1-HMGA1-GFP line and the control line (WT) were immunoprecipitated by anti-GFP beads and western-blotted by anti-GFP or specific antibodies against GH1-HMGA1 or GH1-HMGA2. Different conditions were tested (with or without benzonase and/or formaldehyde). **(H)** Circos plot showing density of peaks of GH1-HMGA1 (a), GH1-HMGA2-Myc (b), GH1-HMGA3 (c) and TE enriched by H3K9me2 (d) mapped on Col-CEN reference genome. **(I)** Venn diagram showing overlap between the identified peaks in the two biological replicates of ChIP-seq targeting GH1-HMGA1 in WT. **(J)** Number of peaks, genes, and transposons identified for each ChiP-seq. **(K)** Heatmap of the three GH1-HMGA proteins enrichment at regions targeted by different combinations of GH1-HMGA.

**Supplementary Figure 2.**
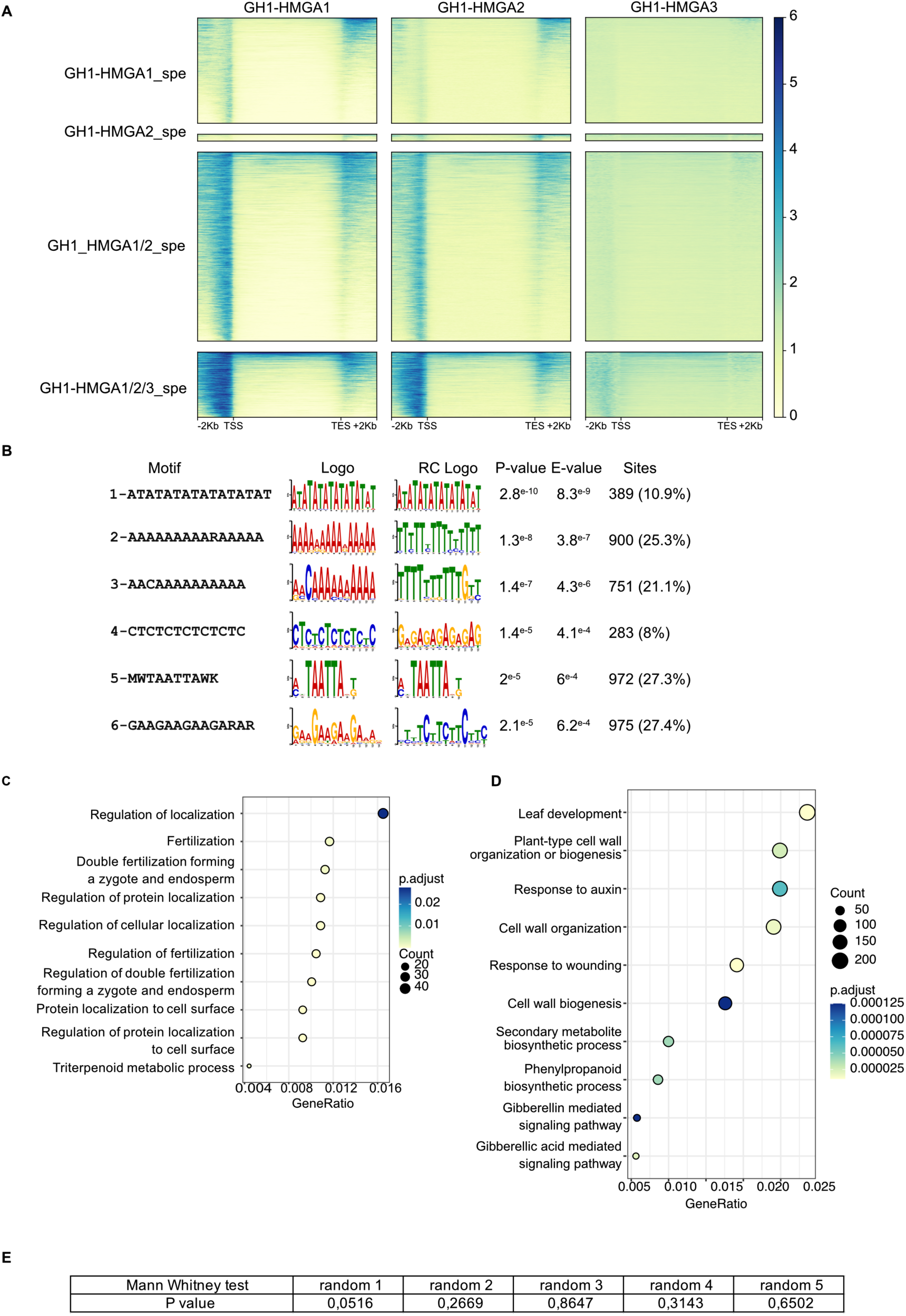
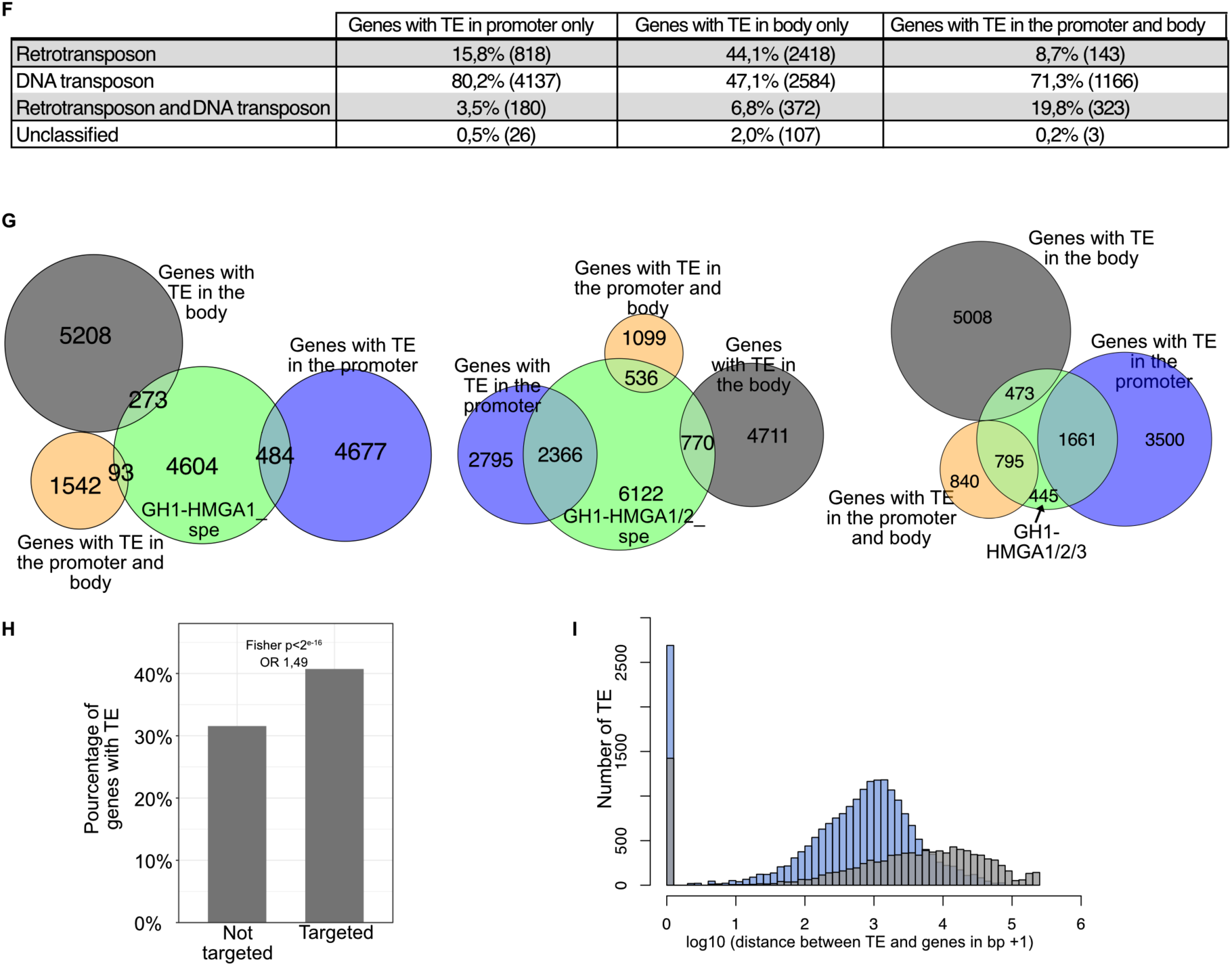
**(A)** Heatmap of the three GH1-HMGA proteins at the genes targeted by different combinations of GH1-HMGA. **(B)** The most abundant DNA sequence motifs identified by XSTREME of peaks in genes **(C)** Gene ontology (GO) term enrichment analysis of genes shared between GH1-HMGA1, GH1-HMGA2 and GH1-HMGA3 performed using clusterProfiler ^40^. **(D)** Gene ontology (GO) term enrichment analysis of genes shared between GH1-HMGA1 and GH1-HMGA2 performed using clusterProfiler. (**E**) Results of five Mann Whitney tests carried out to compare transcription levels between the *gh1-hmga12* mutants and the wild-type (WT) of 500 randomly selected genes (related to Figure 2D). **(F)** The table shows the percentage of *Arabidopsis thaliana* genes containing at least one TE in the promoter region, within the gene body, or in both regions, relative to all annotated genes in the genome. Percentages are further stratified according to TE class: retrotransposons only, DNA transposons only, combined retrotransposons and DNA transposons, and unclassified TEs. **(G)** Euler diagrams illustrate the overlap between different categories of GH1-HMGA target genes (genes specifically bound by GH1-HMGA1, genes commonly bound by GH1-HMGA1 and GH1-HMGA2, and genes bound by GH1-HMGA1, GH1-HMGA2, and GH1-HMGA3) and the corresponding sets of genes containing TEs. **(H)** Percentage of genes containing a transposon (promoter, body or both that are or are not targets of at least one GH1-HMGA protein. Fisher test (odds ratio = 1.49; IC95% [1.42–1.56]; p < 2.2×10⁻¹⁶). **(I)** Distribution of distances between a transposable element (TE) and the nearest gene. GH1-HMGA target TEs are shown in blue and non-target TEs in grey. Distances equal to 0 correspond to TEs overlapping gene coordinate. Wilcoxon rank sum test with continuity correction (p-value < 2.2e-16).

**Supplementary Figure 3.**
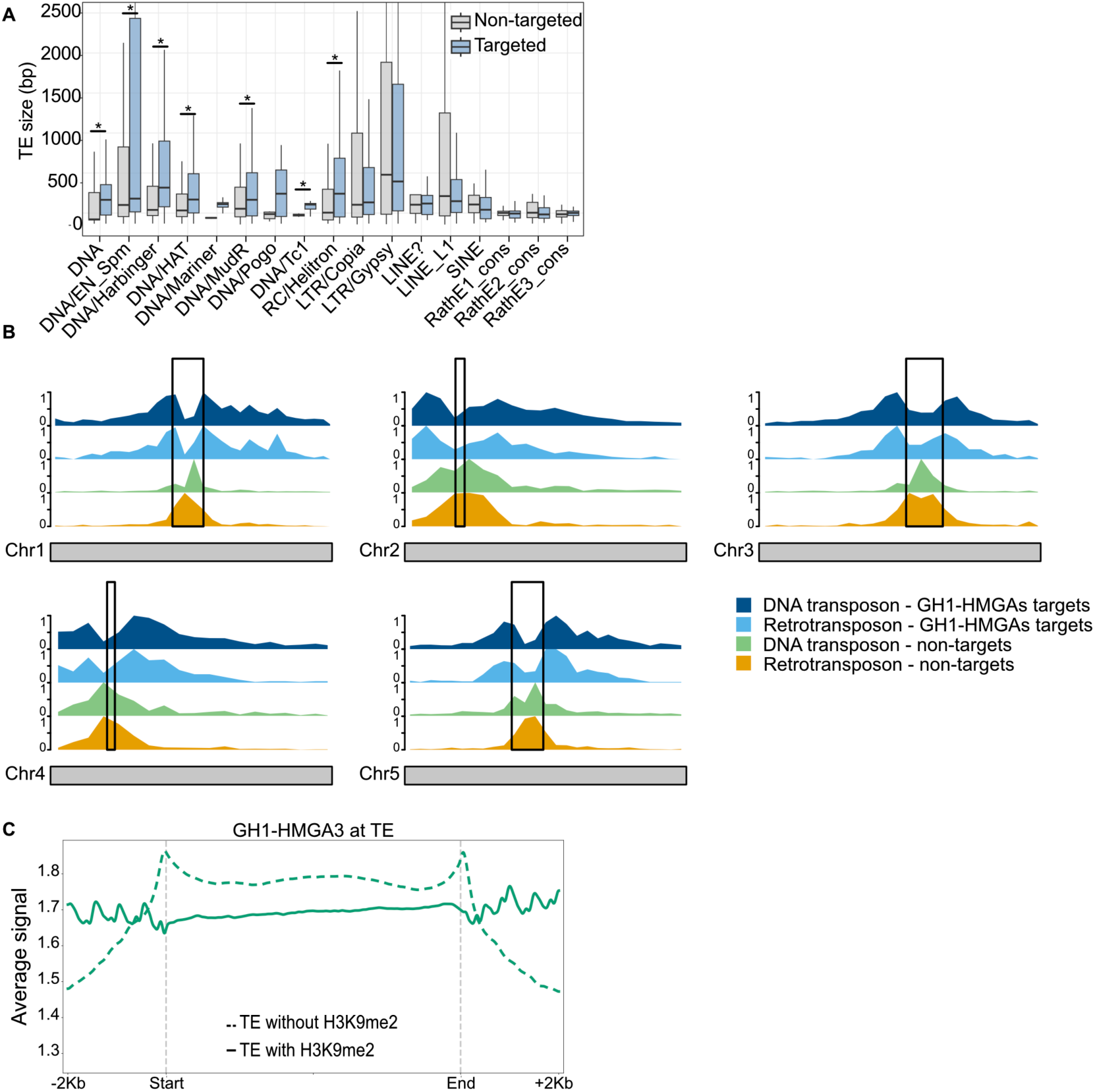
**(A)** Distribution of TE lengths (in base pairs) across TE subfamilies. Boxplots show the distribution of TE lengths within each subfamily for GH1-HMGA target TEs (blue) and non-target TEs (grey). In each boxplot, the central line represents the median, the box boundaries indicate the first and third quartiles (Q1 and Q3), and whiskers representing data spread excluding outliers. Only TE lengths between 0 and 2500 bp are shown (zoomed-in view). Wilcoxon tests were performed to compare the lengths of targeted and non-targeted TEs within each TE subfamily, and p-values were adjusted using the Benjamini–Hochberg method (adjusted p < 0.05). **(B)** Density distribution of transposable elements along the five chromosomes of *Arabidopsis thaliana*. Four density tracks are displayed for each chromosome: DNA transposon targeted (dark blue) and non-targeted (green), retrotransposon targeted (light blue) and non-targeted (orange) by all three GH1-HMGAs. Black boxes indicate centromeric regions as defined by CENH3 enrichment ^41^. **(C)** Metagene plot showing ChIP-seq signal profile of GH1-HMGA3 proteins at TEs enriched or not by H3K9me2.

**Supplementary figure 4.**
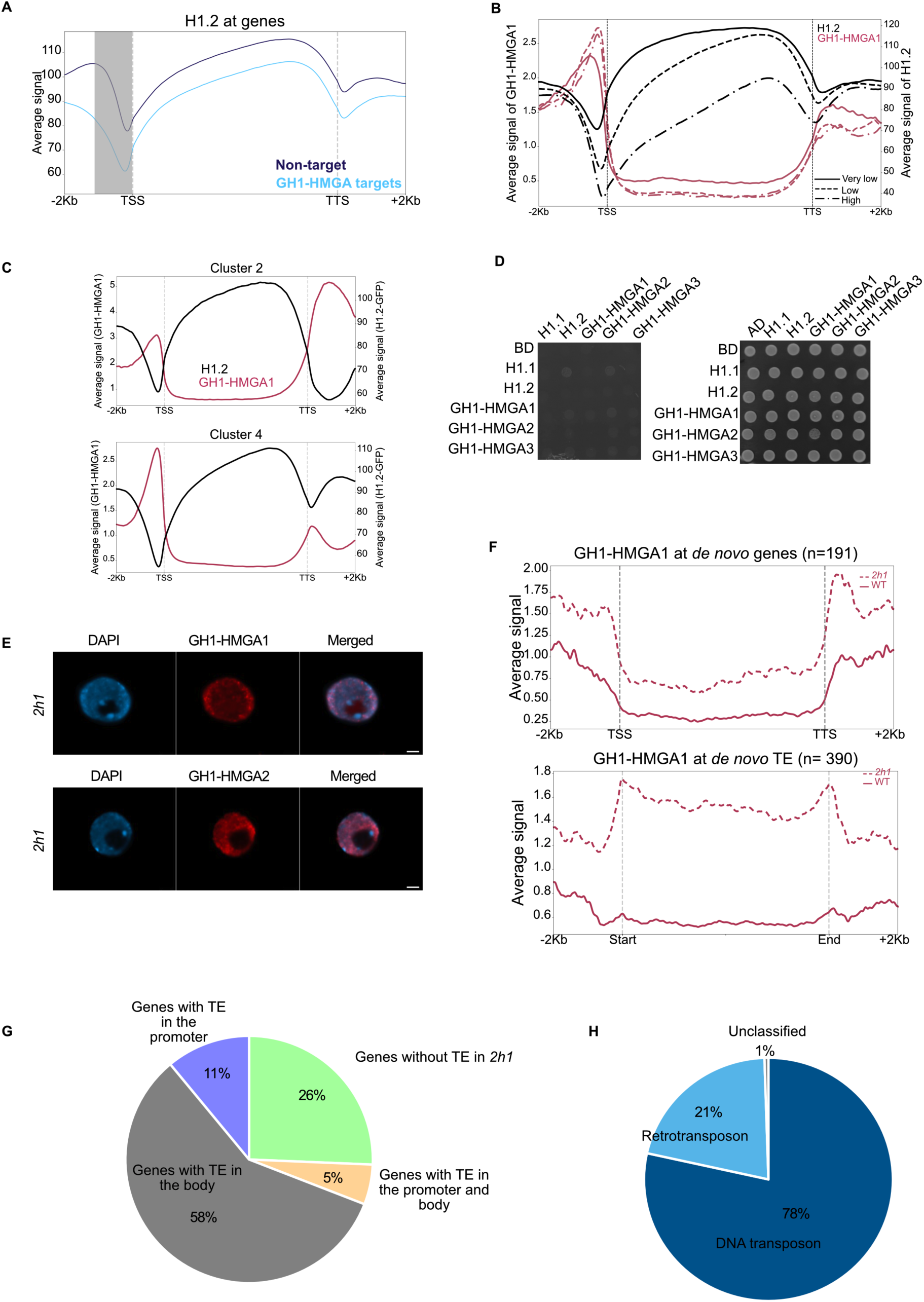
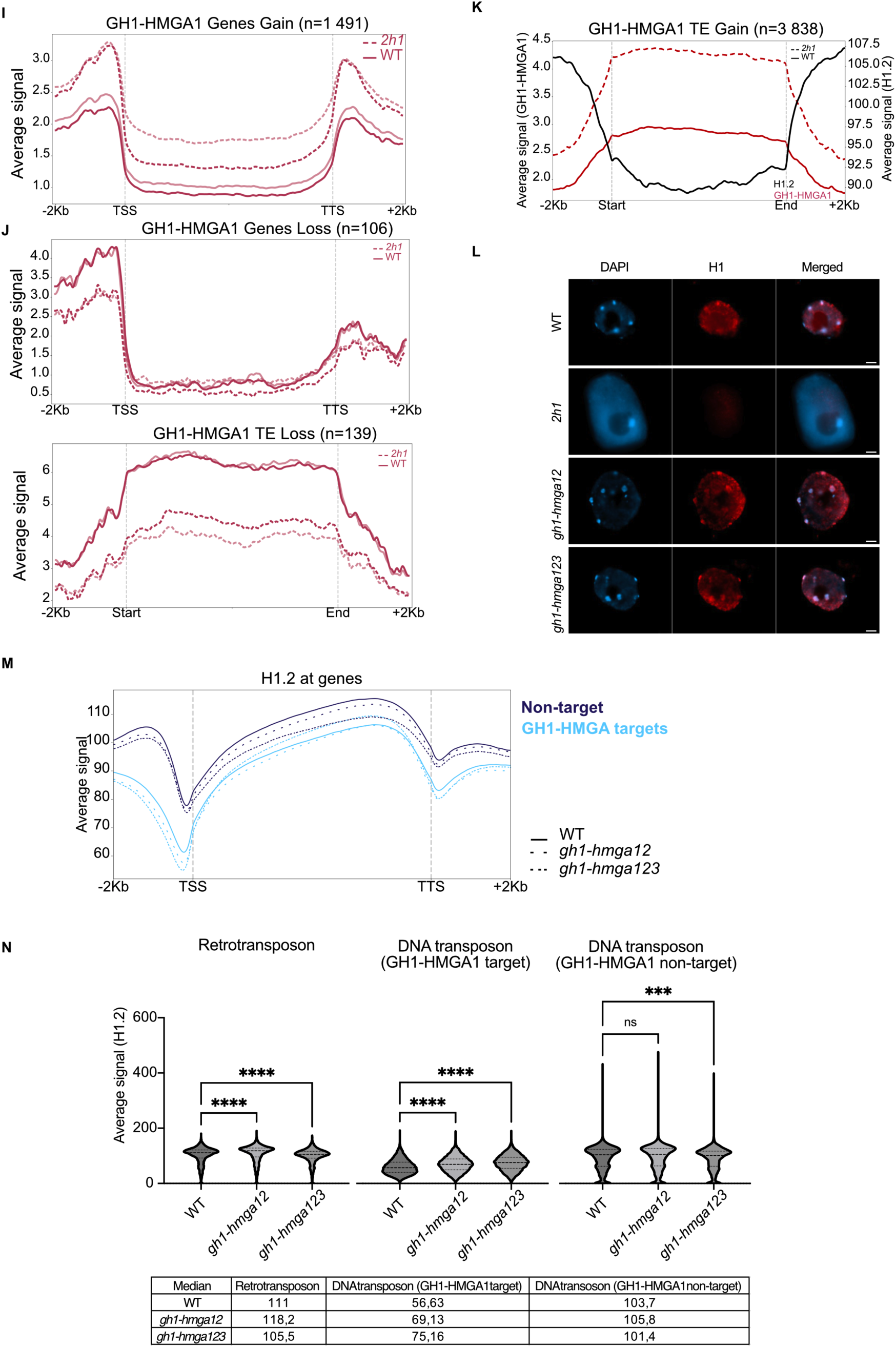
(**A**) Metagene plot showing H1.2-GFP ChIP-seq signal ^17^ over GH1-HMGA1 target genes (n=18723) and over genes not targeted by GH1-HMGA1 (n=14600). The grey shaded area corresponds to the GH1-HMGA1 enrichment region. (**B**) Metagene plots showing GH1-HMGA1 and H1.2-GFP enrichment ^17^ across genes stratified by expression level (very low (n= 3 215), low (n=9 836), high (n=1 771)). (**C**) Metagene plots showing GH1-HMGA1 and H1.2 ChIP-seq enrichment for genes clusters 2 and 4 from Figure 4B. (**D**) Interaction among the three GH1-HMGAs, H1.1 and H1.2 probed in the Y2H system. Growth on selective medium lacking histidine and adenine reveals interaction between the two proteins tested (left panel) and on synthetic medium lacking leucine and tryptophan, selecting for the presence of the bait and prey vectors (right panel). Horizontal, translational fusions with the Gal4-Activation domain (AD); vertical, translational fusion with the Gal4-DNA-binding domain (BD). BD indicate the empty vector. (**E**) Immunostaining of nuclei from 7-days old seedlings using GH1-HMGA1, GH1-HMGA2 antibodies (red) in *2h1* mutants. DNA was counterstained with DAPI (blue, left panels). Merged images of antibody signal and DAPI are shown on the right. Scale bar: 2 µm. (**F**) Metaprofile showing GH1-HMGA1 ChIP-seq signals in wild-type (WT) and *2h1* mutant backgrounds over 191 genes (top) and 390 transposable elements (bottom) *de novo* targeted by GH1-HMGA1 in *2h1*. (**G**) Proportion of new GH1-HMGA1 target genes in the *2h1* mutant according to the presence of TEs in their promoter or body. (**H**) Proportion of new GH1-HMGA1 target TEs in the *2h1* mutant according to transposon class. (**I**) Metaprofile showing GH1-HMGA1 ChIP-seq signal in wild-type (WT) and *2h1* mutant backgrounds over genes displaying significant gain of signal in GH1-HMGA1. Two biological replicates are displayed with different shades of the same colour. (**J**) Metaprofiles showing GH1-HMGA1 ChIP-seq signal in wild-type (WT) and *2h1* mutant backgrounds over genes and transposable elements displaying significantly lower enrichment in GH1-HMGA1 in *2h1* compared to WT as determined by DESeq2 (adjusted p-value < 0.05). Two biological replicates are displayed with different shades of the same colour. (**K**) Metagene profile showing H1.2 and GH1-HMGA1 ChIP-seq signals in WT and *2h1* mutant background over TE displaying significant higher enrichment in GH1-HMGA1 in *2h1* compared to WT as determined by DESeq2 (n=3838). **(L)** Immunostaining of nuclei from 7-days old seedlings using an H1 antibody (red) in *2h1, gh1-hmga12* and *gh1-hmga123* mutants. DNA was counterstained with DAPI (blue, left panels). Merged images of antibody signal and DAPI are shown on the right. Scale bar: 2 µm. (**M**) Metagene plot showing mean H1 ChIP-seq signal from 2 biological replicates over GH1-HMGA1 target genes (n=18723) and over genes not targeted by GH1-HMGA1 (n=14600) in *gh1-hmga12*, *gh1-hmga123* and WT seedlings. (**N**) Violin plot of ChIP-seq signal and median of average signal of H1 in WT, *gh1-hmga12* and *gh1-hmga123* at retrotransposons and DNA transposons targeted or not by GH1-HMGA1. Dunn’s multiple comparisons test (adjusted p-value <0.0001). The values of the medians are indicated in the table.

**Supplementary figure 5:**
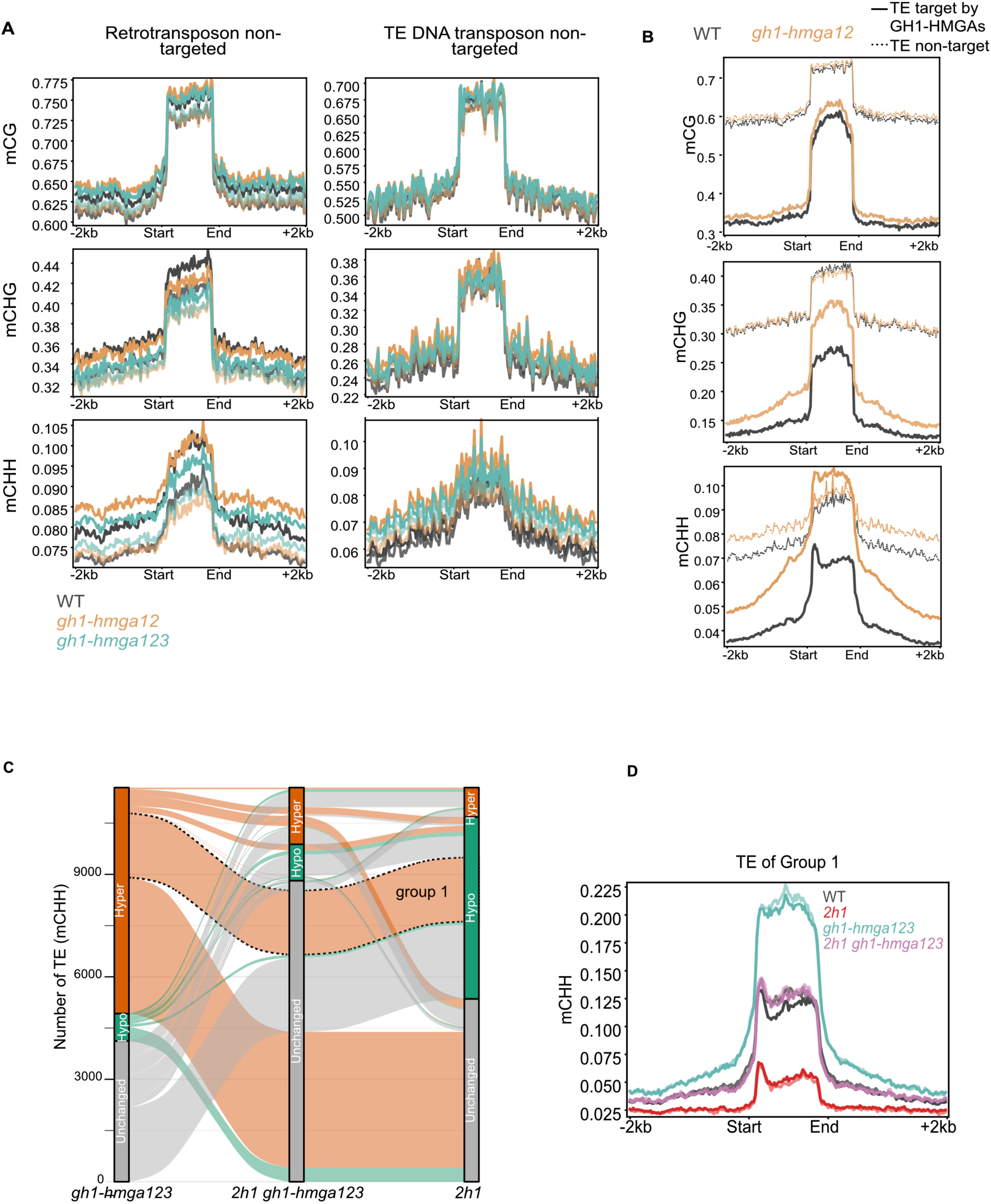
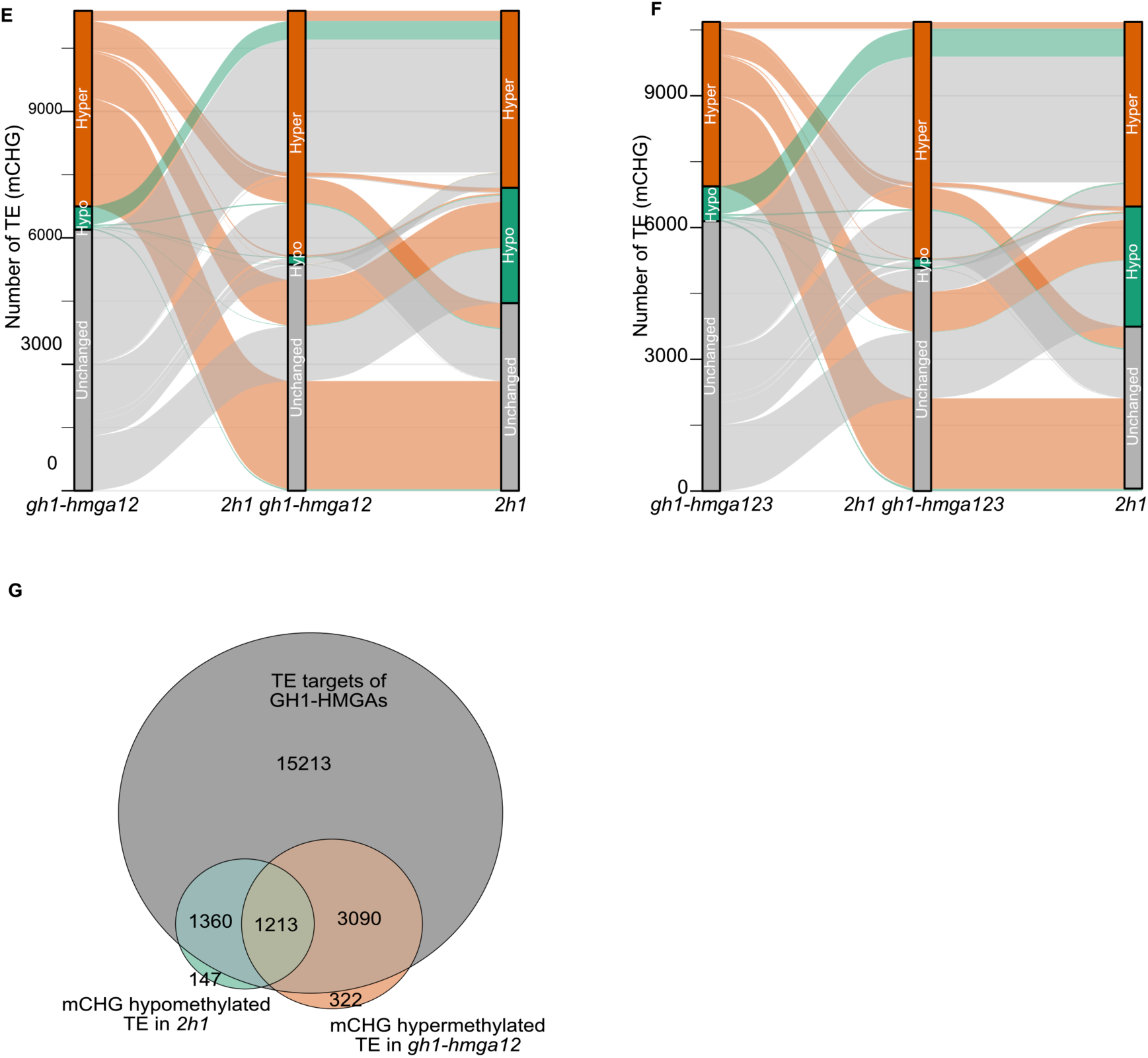
Supplemental information related to Figure 5. **(A)** Metaprofile showing Bisulfite-seq signal profile (mCG, mCHG, mCHH) across retrotransposons and DNA transposons non-targeted by GH1-HMGAs for WT, *gh1-hmga12* and *gh1-hmga123*. Two biological replicates are displayed in the same color with different shades. **(B)** Metaprofile showing Bisulfite-seq signal profile (mCG, mCHG, mCHH) across TEs targeted and non-targeted in WT and *gh1-hmga12*. **(C)** Alluvial diagram of CHH methylation changes in TEs in *2h1*, *gh1-hmga123*, and *2h1 gh1-hmga123* mutants. Hypermethylated, hypomethylated, and unchanged TEs relative to WT are shown in orange, green, and grey, respectively. TEs hypermethylated in *gh1-hmga123*, hypomethylated in *2h1*, and unchanged in *2h1 gh1-hmga123* relative to WT are outlined with a dotted line. **(D)** Metaprofile of Bisulfite-seq mCHH signal across the TE group 1 in (C). **(E)** Alluvial diagram of CHG methylation changes in TEs in *2h1*, *gh1-hmga12*, and *2h1 gh1-hmga12* mutants. Hypermethylated, hypomethylated, and unchanged TEs relative to WT are shown in orange, green, and grey, respectively. TEs hypermethylated in *gh1-hmga12*, hypomethylated in *2h1*, and unchanged in *2h1 gh1-hmga12* relative to WT are outlined with a dotted line. **(F)** Alluvial diagram of CHG methylation changes in TEs in *2h1*, *gh1-hmga123*, and *2h1 gh1-hmga123* mutants. Hypermethylated, hypomethylated, and unchanged TEs relative to WT are shown in orange, green, and grey, respectively. TEs hypermethylated in *gh1-hmga123*, hypomethylated in *2h1*, and unchanged in *2h1 gh1-hmga123* relative to WT are outlined with a dotted line. **(G)** Euler diagram showing overlap between GH1-HMGA target TEs and the TE either CHG hypomethylated in *2h1* or CHH hypermethylated in *gh1-hmga12*.

**Supplementary figure 6:**
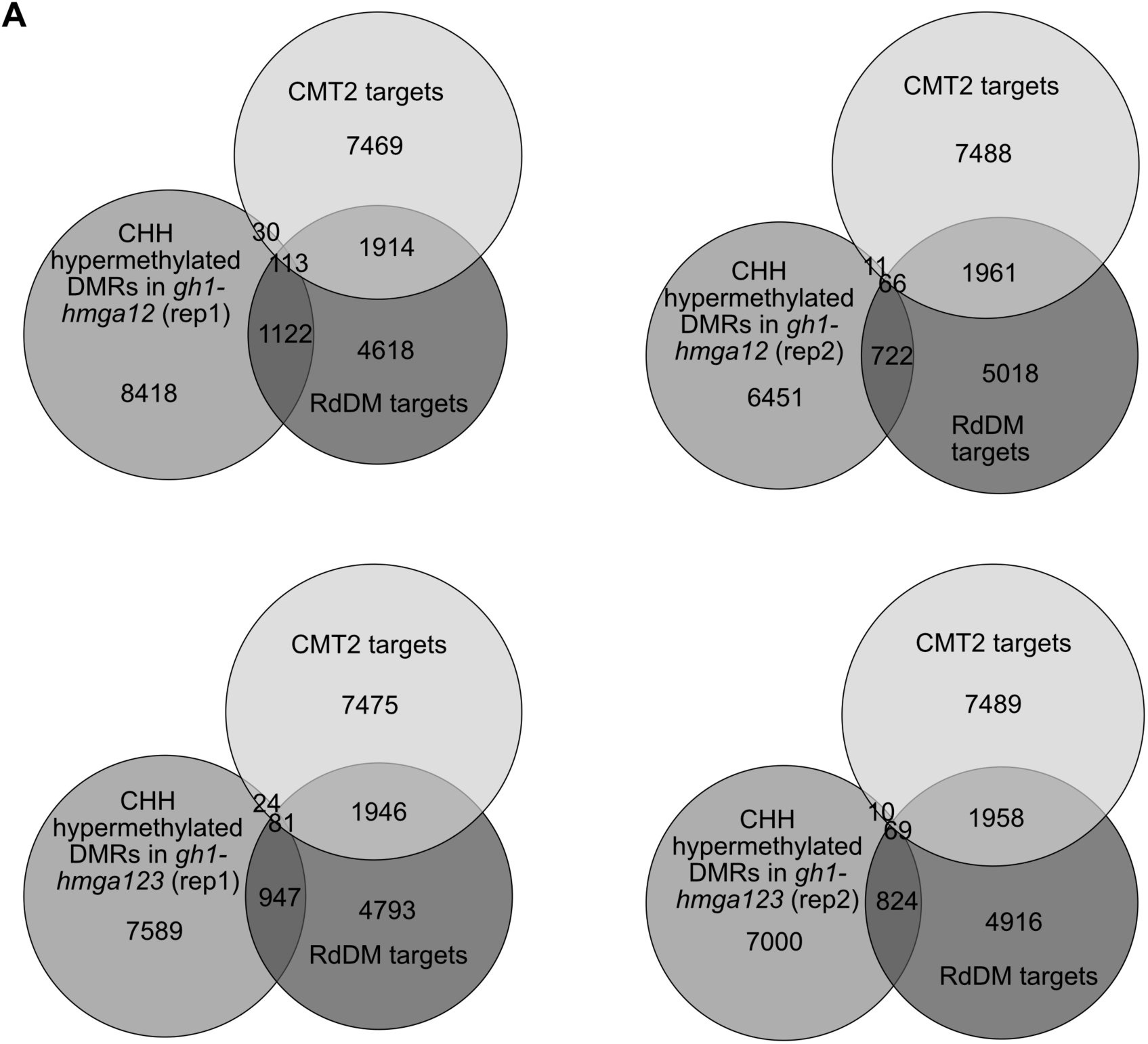
Supplemental information related to Figure 6. **(A)** Euler diagrams showing the overlap between CMT2-, RdDM-targeted regions and CHH hypermethylated DMRs identified in *gh1-hmga12* and *gh1-hmga123* mutant replicates.

## Notes

### Competing Interest Statement

The authors have declared no competing interest.

